# Curvature induced patterns: A geometric, analytical approach to understanding a mechanochemical model

**DOI:** 10.1101/2025.04.17.649327

**Authors:** Daphne Nesenberend, Arjen Doelman, Frits Veerman

## Abstract

The exact mechanisms behind many morphogenic processes are still a mystery. Mechanical cues, such as curvature, play an important role when tissue or cell shape is formed. In this work, we derive and analyze a mechanochemical model. This particular spatially one-dimensional model describes the deformation of a tissue- or cell surface over time, which is driven by a morphogen that locally induces curvature. The model consists of two PDEs with periodic boundary conditions; one reaction-diffusion equation for the morphogen and one PDE that describes the dynamics of the curve, derived by taking the *L*^2^-gradient flow of the Helfrich energy. We analyze the possible steady states of this model using geometric singular perturbation theory. It turns out that the strength of interaction between the morphogen and the curvature plays a key role in the type of possible steady state solutions. In the case of weak interaction, the geometry of the slow manifolds allows only for (in space) slowly changing periodic orbits that lay completely on one slow manifold. In the case of strong interaction, there exist multiple front solutions: periodic orbits that jump between different slow manifolds. The singular skeletons of the steady state solutions do not meet the required consistency conditions for the curvature, a priori indicating that the solutions might not be observable. The observability and stability are investigated further using numerical simulation.

## 1 Introduction

Organisms are able to develop complex spatial structures on many different scales. From organs to individual cells, these structures have morphological features to fulfill their specific functions optimally. For example, the human intestines have a tortuous build, evolutionary driven to maximize the total nutrient absorption surface. Zooming in on the intestine wall, one finds cells that have many protrusions, highly deforming the cell membrane and further enlarging the absorption surface [3]. But how do these cells “decide” to form such regular spatial structures? What mechanisms drive these patterns to appear?

To answer these questions, mathematical modeling can be employed. Alan Turing showed that two coupled reaction diffusion equations could display the formation of spatially inhomogeneous patterns [34]. These equations describe two chemical species that diffuse in a spatial domain: a short-range activator and a long-range inhibitor. When the homogeneous background state is unstable with respect to spatial perturbations, patterns are formed. Others have followed this approach to explain different evolutionary processes [8; 38]. However, in several applications [5; 11], it has been difficult to identify a chemical that could play the role of the long-range inhibitor required in Turing’s classical approach [36].

Cells and molecules are also sensitive to mechanical signals [14; 30; 9; 25]. Mechanical cues such as shape, stress, and strain can be considered to serve as an alternative source of regulation [35; 19; 13; 10]. Several different models have been proposed to describe this mechanochemical interaction, focusing on the elastic properties of the underlying surface [26; 24; 23; 39; 37] or describing the effective interaction between the local curvature of a thin flexible membrane with the morphogen [10]. In this paper, we follow the approach of Mercker et al., in which a mechanochemical model is derived and studied, where mechanical constraints of the surface take over the role of the inhibitor [29; 36]. The model describes an activating molecule (the morphogen) that diffuses on a surface and locally induces the curvature of the surface. Continuity and compressibility constraints of the surface induce a long-range inhibitory effect on the surface curvature. Numerical simulations show that mechanochemical models can exhibit a wide range of patterns [28; 10; 31].

In this work, we derive a one-dimensional mechanochemical model that describes the interaction between a deforming planar curve Γ ⊂ ℝ ^2^ and a curvature-inducing morphogen Φ that is localized on Γ. The model equations will be derived from first principles, using a similar approach as Mercker et al. [29]. Our model is based on a modified Helfrich energy [16],

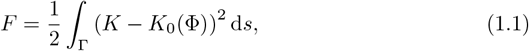

where *K* is the local curvature of Γ, *K*_0_ the preferred local curvature dependent on morphogen concentration Φ, and *s* the arc length coordinate along the curve Γ. We make the additional assumption that the curve Γ can for all time *t ≥* 0 be written as the graph over *x* ∈ [0, *L*] of a function *y* = *h*(*x, t*) in a fixed Cartesian (*x, y*)-coordinate system. From this assumption, it follows that the curvature *K* can be expressed in terms of *h* as

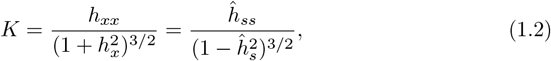

where

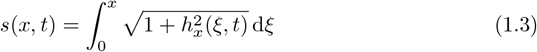

is the arc length coordinate of the curve Γ, and *ĥ*(*s*(*x, t*), *t*) = *h*(*x, t*). In Section 2 we take the *L*^2^-gradient flow of the modified Helfrich energy functional (1.1) to derive the evolution equation that minimizes this energy over time. Together with some other assumptions, we obtain the following system of governing partial differential equations:

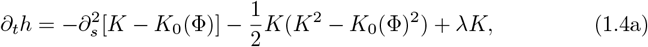

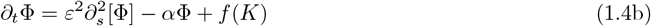

for the evolution of the curve (1.4a) and of the morphogen (1.4b), respectively. Here, *λ* is the Lagrange multiplier that incorporates the global arc length constraint

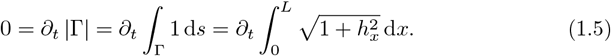

We assume that the morphogen Φ is diffusing significantly more slowly than the speed with which the surface adjusts to its preferred local curvature, encoded by 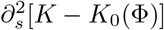 in (1.4a), leading to the scaling assumption 0 *< ε* ≪1. The morphogen is constantly degraded with rate *α* and produced with rate *f* (*K*). A complete derivation of (1.4) and an overview of all assumptions underlying this model can be found in Section 2.

In Section 3 we perform a steady state analysis of (1.4), to investigate and establish the existence of localized and spatially periodic patterns. We analytically investigate the steady states of system (1.4) using geometric singular perturbation theory (GSPT) [20; 22]. GSPT uses the fact that the small diffusion rate *ε* induces a spatial scale separation. This scale separation can be employed to analyze the existence and structure of stationary solutions to (1.4), which are solutions to the ODE system (3.5). From the geometric properties of the curve, i.e. the graph constraint and the periodic boundary conditions, we derive a set of geometric consistency conditions that a solution needs to comply with to be observable.

The dynamics of system (3.5) can be investigated by analyzing the lower-dimensional subsystems (3.51) and (3.56)/(3.53). The dynamics of these subsystems can be combined using geometric arguments to construct stationary pattern solutions to (1.4). Using these techniques, we identify a pattern bifurcation parameter *A* (3.7) that is a measure for the strength of interaction between the morphogen and the curvature. For weak interaction (*A <* 1), we find patterns that completely lie on the slow manifold – i.e., that do not exhibit any sharp spikes or fronts. For strong interaction (*A >* 1) we prove the existence of solutions that jump between slow manifolds for *ε* sufficiently small, yielding multiple front-type structures in the associated pattern. Furthermore, we investigate the observability of these patterns by checking the geometric consistency of the curve represented by the steady state orbits. Here we find that the singular skeleton of the periodic solutions cannot comply with the geometric consistency conditions of the curve. However, we argue that there might be solutions close to singular skeleton but sufficiently distant such that they can comply with the geometric consistency conditions i.e. *ε* small, but not too small.

The analytical approach using GSPT allows us to extend our understanding beyond “close to equilibrium” patterns such that we can explore a range of possible patterns, including more exotic solutions like sharp spikes and fronts, which can be classified as “far from equilibrium” patterns. GSPT-based techniques have been used to analyze patterns in a wide range of pattern-forming systems [15].

In Section 4, we complement our analytical findings on the existence of stationary patterns with a numerical simulation of the full PDEs (1.4), to investigate the stability of the stationary patterns whose existence we established in Section 3. We find that this system exhibits a range of periodic patterns. In Section 5, we discuss our results and provide suggestions for future research.

## 2 Model derivation

In this section we derive a mechanochemical model on a curve. This model describes the evolution of a curve and a morphogen diffusing on that curve. There is an interaction between the local curvature and the morphogen: the curvature is inducing morphogen production, and the morphogen is inducing curvature in turn. Assuming a fixed curve length, there is a long range inhibiting effect of the curve on itself. Together, these processes make pattern formation possible. We will discuss the assumptions underlying the model and derive the evolution equation of the curve from first principles using a similar approach as Mercker et al. [29].

### 2.1 Setup and context

We consider a planar curve Γ ⊂ ℝ^2^. Our first assumption is that Γ can be written as a graph over *x*∈ [0, *L*] of a function *y* = *h*(*x, t*) in a fixed Cartesian (*x, y*)-coordinate system for all time *t ≥* 0. The local curvature *K* of the curve Γ is the main driver of the evolution of the shape of the curve. In the given graph representation, we know that

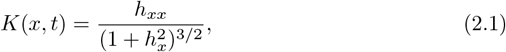

where *h*_*x*_ = *∂*_*x*_*h*(*x, t*) and 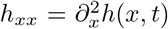 denote the partial spatial derivatives of *h* [33]. Furthermore, the metric tensor *g* of the curve Γ is given in terms of *h* as

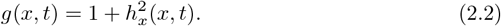

We introduce the arc length coordinate

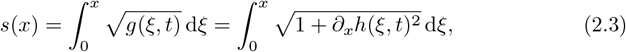

and introduce the function *ĥ* which describes Γ as a graph in the arc length coordinate, i.e.

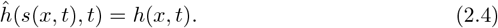

Note that by applying the chain rule, we can express the partial derivative to *x* in terms of *s* as

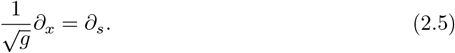

We can derive that in the arc length coordinate *s*, the curvature can be expressed as

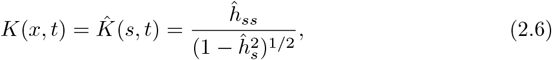

since we can derive from (2.5) that

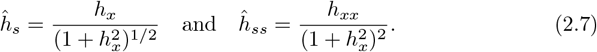

### 2.2 Curve dynamics

The starting point of our model is the assumption that the curve Γ tends to minimize its modified Helfrich energy

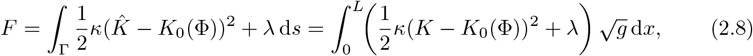

where the preferred local curvature *K*_0_ depends on the local morphogen concentration Φ, that we later explicitly choose. For simplicity, we assume that the bending modulus *κ* is independent of Φ; however, the analysis in this paper can be straightforwardly extended to the case where *κ* depends on Φ in a nontrivial manner.

The Lagrange multiplier *λ* constrains the curve to keep a constant fixed length,

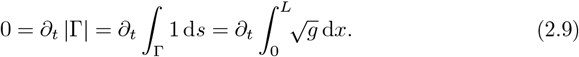

For future reference, we denote the length of Γ by

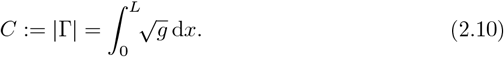

The total energy *F* becomes larger (less favorable) when the local curvature is further diverted from the preferred local curvature *K*_0_. Note that, in contrast to [29], we do not assume a local incompressibility constraint 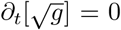, since that would induce *∂*_*t*_[*h*_*x*_] = 0, which would prohibit any nontrivial curve deformations.

We assume that the evolution of Γ is governed by the *L*^2^-gradient flow

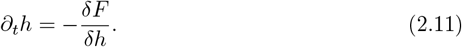

We calculate the variational derivative 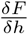 by substituting *h* → *h* + *ϵp*, with *p*(0, *t*) = *p*(*L, t*) = *p*_*x*_(0, *t*) = *p*_*x*_(*L, t*) = 0 and expanding the resulting expression to first order in *ϵ*. We find

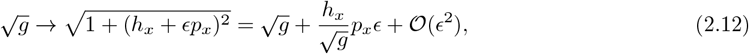

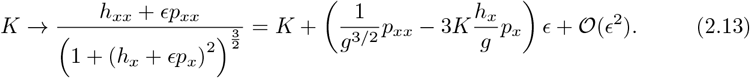

Substituting these expansions in *F* (2.8), we obtain

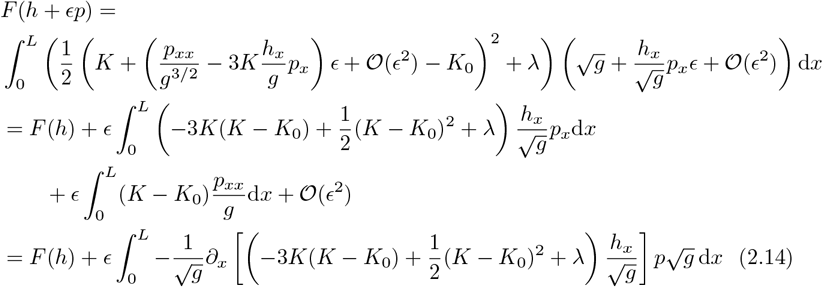

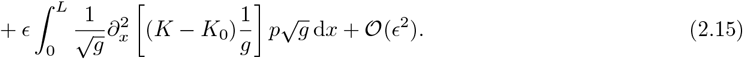

We can simplify the above by expressing the partial derivatives in terms of the arc length *s*, using (2.5). We first observe that

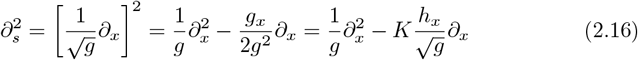

and

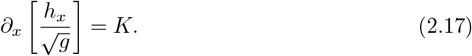

Define *u* := *K* − *K*_0_. We simplify the functional part of the integrand of (2.14) as

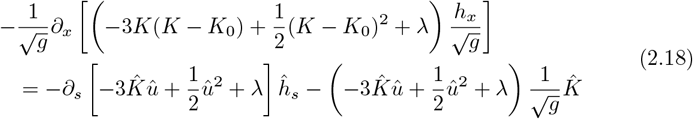

and of (2.15) as

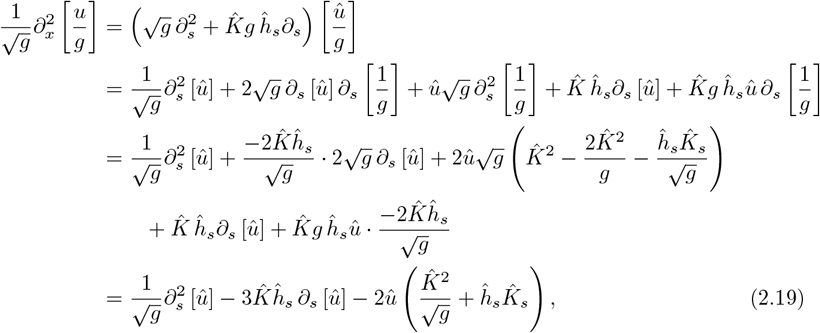

where we have used 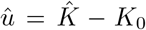 to highlight the coordinate change from *x* to *s*. Combining (2.18) and (2.19) in (2.14)/(2.15), we calculate the difference quotient

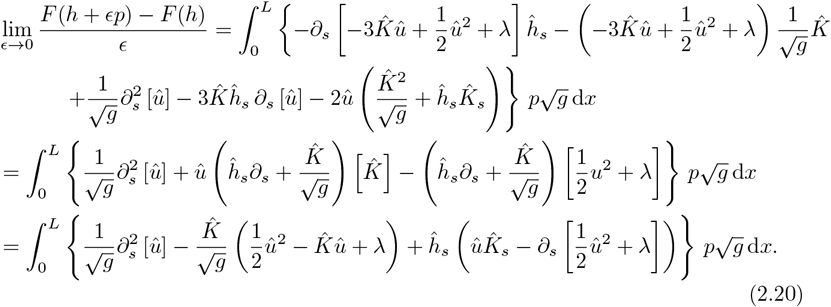

From the relation

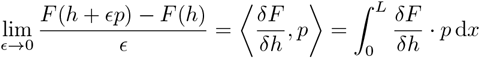

we derive that

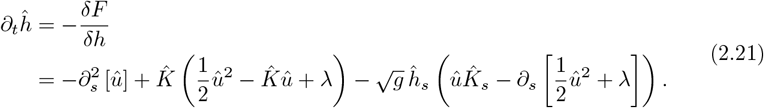

Since the Darboux frame of the curve (*s, ĥ*(*s, t*)) is spanned by (1, *ĥ*_*s*_) and (−*ĥ*_*s*_, 1) and *∂*_*t*_ (*s, ĥ*(*s, t*)) = (0, *∂*_*t*_*ĥ*), the last term 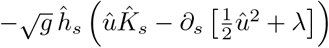 in equation (2.21) quantifies the variation of the curve in the tangential direction; see also [29]. For a closed curve (or, more general, for a surface without boundary) this term can be associated to a reparametrization of the curve [2] (see also [4; 12]). Since we are interested in quantities that are invariant under curve reparametrizations, and since we will be imposing periodic boundary conditions (cf. Section 2.4), we omit this tangential variation term. We obtain (dropping the hats)

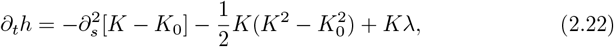

as the evolution equation for the curve dynamics.

### 2.3 Morphogen dynamics

We assume there to be a linear relationship between the local preferred curvature and the morphogen concentration,

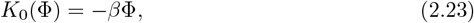

to have a simple expression where morphogen induces negative curvature. However, one can tailor *K*_0_ to express specific biological implementations. We assume that the morphogen concentration on the curve

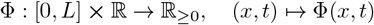

is governed by a the reaction diffusion equation

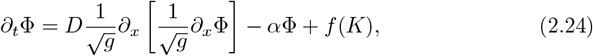

which can be written in terms of *s* as

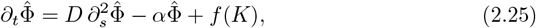

where 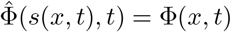. Here, *D* is the diffusion coefficient of the morphogen, *α* is the degradation of the morphogen and *f* (*K*) is the curvature dependent morphogen production. Note that in (2.24) we use the Laplace-Beltrami operator instead of the standard Laplacian, since the morphogen is restricted to the curve Γ. We assume that the diffusion of the morphogen is slow compared to the relaxation rate of the curve: hence, we write *D* = *ε*^2^, where 0 *< ε* ≪ 1.

In this paper, we do not have a specific biological system in mind, so we cannot use that to deduce an interaction function *f* (*K*). For now, we only (exclusively) want negative curvature to induce morphogen production, so *f* (*K*) = 0 when *K >* 0 and *f* (*K*) *>* 0 when *K <* 0. Furthermore, we want the production to be uniformly bounded. We choose *f* to be a continuous, piecewise differentiable function,

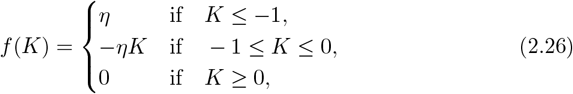

that simplifies the analysis to be performed on the system. Note that this function has similar properties to Hill functions, often used in regulatory networks [32].

#### 2.3.1 The piecewise smooth nature of *f*

The fact that *f* (*K*) (2.26) is not continuously differentiable introduces technical difficulties in the existence analysis in Section 3. To circumvent these issues, we replace the piecewise smooth function *f* (*K*) (2.26) in the statements of our theorems by a smooth variant 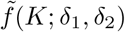 that depends on two auxiliary small parameters *δ*_1,2_, such that 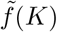is uniformly *𝒪* (*δ*_1_, *δ*_2_) close to the original *f* (*K*) for fixed small *δ*_1_ and *δ*_2_. Here, *δ*_1_ is used to define a small interval around the corner points *K* = −1 and *K* = 0, and *δ*_2_ is used to control the slope deviation in between *K* = − 1 and *K* = 0. An example of a *C*^2^ extension of *f* (*K*) would be

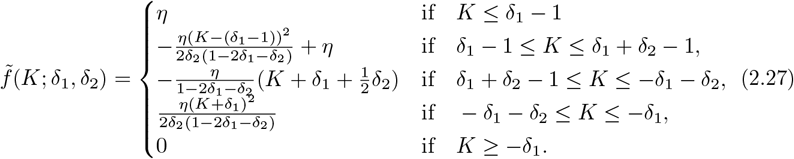

Other choices of sufficiently smooth regularizations of *f* (*K*) are also possible. For the analysis of this work, we do not choose a particular regularization; instead, we work with the expression (2.26) and prove existence of stationary periodic orbits in system (1.4) for any smooth regularization 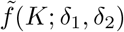 for sufficiently small fixed *δ*_1,2_. The defining properties of such a regularization 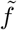 are

- 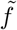 is smooth in *K, δ*_1_ and *δ*_2_, and is monotonically decreasing in *K*;
- 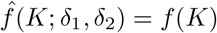 for *K* ≤ *δ*_1_ − 1 and for *K ≥* −*δ*_1_;
- 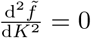 for *δ* + *δ*_2_ − 1 ≤ *K* ≤ −*δ*_1_ − *δ*_2_;
- 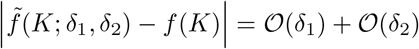 uniformly in *K*.

### 2.4 Boundary conditions

The assumption of a global graph representation of Γ allows for vertical translations of the curve; indeed, system (1.4) is invariant under *h*(*x, t*) → *h*(*x, t*) + *h*_0_ for fixed *h*_0_. Hence, for the full PDE simulations discussed in Section 4, we can fix *h*(0, *t*) = 0.3 without loss of generality.

The global graph representation of Γ does not a priori provide natural boundary conditions on *h*_*x*_ and/or *h*_*t*_. However, we can imagine our graph to be a small section of a simple closed curve, which is uniformly close to a circle with a large radius. In this extended setting, studying periodic patterns on this large circle is equivalent to choosing *L* such that an integer multiple of *L* is equal to 2*πR*, and posing spatially periodic boundary conditions on *h*(*x, t*). With this extended embedding in mind, we set

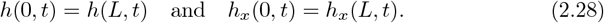

As the curve evolution equation (1.4a) contains second order spatial derivatives of the curvature (and is therefore of fourth order in *h*), we extend the spatially periodic boundary conditions to *K*, and set

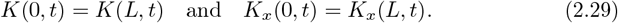

Following the same reasoning that led to the spatially periodic boundary conditions on *h*, we impose on the morphogen

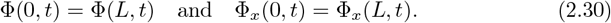

## 3 Spatially periodic patterns

In this section we analytically determine the existence of stationary, spatially periodic solutions to system (1.4). That is, we consider time independent solutions to (1.4), and thus set *∂*_*t*_*h* = *∂*_*t*_Φ = 0. As we will be using *s* as independent variable throughout Section 3, we will omit the hat-notation employed in Section 2, for clarity of notation.

We introduce

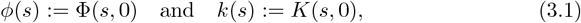

and define

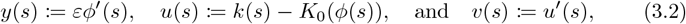

which allow us to study stationary solutions to the PDE system (1.4) as solutions to the following system of ODEs:

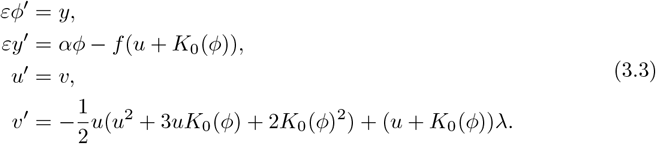

Observe that system (3.3) is dependent on curvature *K*, but not on height *h*. This reduces the dimensionality of the problem from six to four. Furthermore, note that the assumption 0 *< ε* ≪ 1, indicating a significant difference between the diffusivities of *u* and *ϕ*, makes system (3.3) singularly perturbed. This singularly perturbed structure induces a scale separation in the flow of (3.3), and allows for the use of geometric singular perturbation theory (GSPT) to analytically investigate the existence of several types of orbits in (3.3). The specific choices for the interaction functions *f* (*K*) and *K*_0_(*ϕ*) can be found in (2.26) and (2.23). To slightly simplify system (3.3), we rescale the asymptotically small parameter 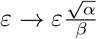, and introduce

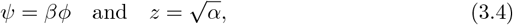

to obtain

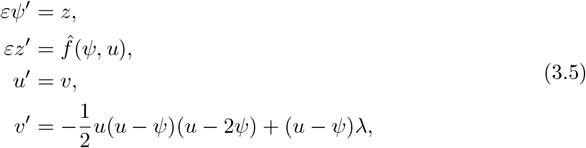

where (cf. (2.26))

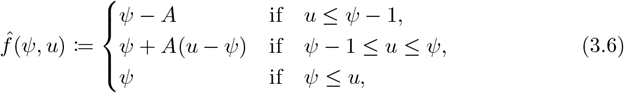

with

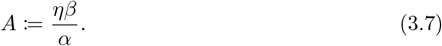

Note that 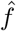 (3.6) is piecewise smooth. When appropriate, we will replace 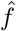 by a suitable smooth regularization, see Section 2.3.1 for details. As a preparation for our upcoming analysis, we rescale the spatial variable with *ε*, and introduce 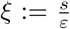. Denoting 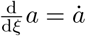, system (3.5) becomes

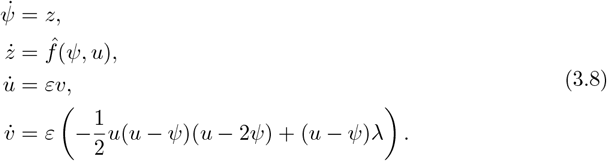

We refer to (3.5) as the slow system and to (3.8) as the fast system.

The key parameter that determines the behavior of solutions to (3.5)/(3.8) is *A* (3.7), which quantifies the interaction strength between the morphogen and the curvature.

In the following subsections, we analyze system (3.3), using GSPT. First, we obtain normally hyperbolic critical manifolds, whose geometry depends on the bifurcation parameter *A* (3.7). Then we determine several types of pattern solutions for two fundamentally different regimes of parameter *A* and check their viability as solutions to the PDE (1.4).

### 3.1 Geometric consistency conditions for an evolving curve

All steady states to the PDE system (1.4) are orbits in the phase space of the ODE system (3.3). However, not all these orbits are observable solutions to the systems of PDEs (1.4). For an orbit to be an observable solution to the PDE system (1.4), three conditions need to be met:

1. the curvature *K*(*s*) yields a spatial curve that can be written as a graph *h*(*x*),
2. this graph agrees with the boundary conditions discussed in Section 2.4,
3. the corresponding steady state is stable (in the temporal sense) as a solution to (1.4).

The first two conditions restrict the geometry of the curve associated to a solution *K*(*s*) to (3.3); we will therefore refer to conditions 1 and 2 as the *geometric consistency conditions*. As the geometric consistency conditions directly influence the upcoming existence analysis, we will explore these in more detail below.

The periodic boundary conditions (2.29) imply that *K*(*s*) is *C*-periodic, see (2.10). Hence, let the curvature *K*(*s*) be a periodic orbit with period *C*. Modulo rotation and translation, the curvature uniquely defines a curve Γ. Our goal is to derive necessary conditions on *K*(*s*) such that Γ has a global graph representation *h*(*x*) for 0 ≤*x* ≤*L* with periodic boundary conditions (2.28). We can rewrite the periodic boundary conditions in terms of the arc length coordinate *s* (2.3) as

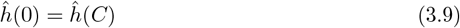

and

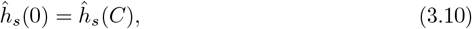

where *ĥ* is defined in (2.4). Assume that Γ has a global graph representation. We use that

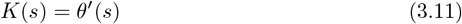

where *θ* is the tangent angle [33]. Hence, by integrating over the curvature, the rotation of the tangent vector with respect to the initial tangent vector, 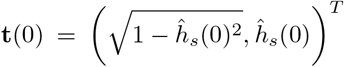, can be derived, see (3.14). Now we use that the tangent vector is equal the derivative of the curve, **t**(*s*) = (*x*_*s*_, *ĥ*_*s*_)^*T*^ [33], to derive that

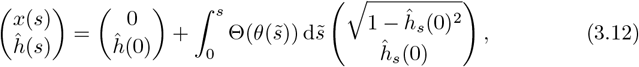

with rotation matrix

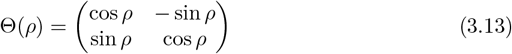

and net tangent vector rotation

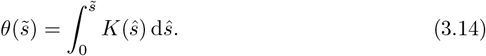

Boundary condition (3.10) yields

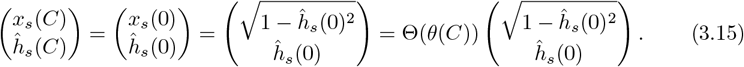

Since Θ(*θ*(*C*)) is the matrix representation of a clockwise rotation over an angle *θ*(*C*), it follows that 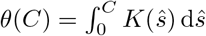 must be a multiple of 2*π*. However, as

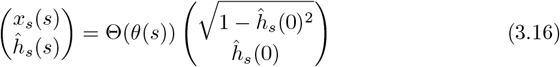

holds for all 0 ≤ *s* ≤ *C*, we can conclude more. Due to the graph representation, it is necessary that *x*(*s*) is strictly increasing, i.e. 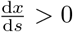 for all 0 ≤ *s* ≤ *C*. Since the vector 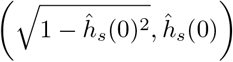 is a unit vector in the positive half plane with angle arcsin *ĥ*_*s*_(0) with respect to the horizontal axis, and Θ(*θ*(*s*)) is the clockwise rotation around the origin over an angle *θ*(*s*), it follows that 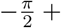 arcsin 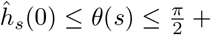 arcsin *ĥ*_*s*_(0) for all 0 ≤ *s* ≤ *C*, so that *(x*_*s*_(*s*), *ĥ*_*s*_(*s*)) remains in the right half plane. We conclude that

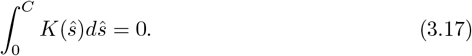

Boundary condition (3.9) yields

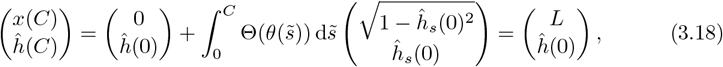

from which it follows that

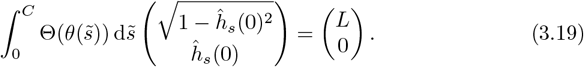

Condition (3.19) provides a nontrivial relation between *L* and *C*, given *ĥ*_*s*_(0).

### 3.2 Phase space analysis and invariant manifolds

Stationary solutions to (1.4) that obey the periodic boundary conditions (2.28)-(2.30) correspond to periodic orbits in system (3.3). To study the structure of the phase space of system (3.3), we start by identifying invariant lower-dimensional manifolds that will turn out to govern the dynamics of (3.3).

We take the singular limit *ε* ↓ 0 in system (3.5) to obtain

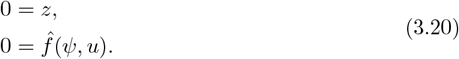

The algebraic equations (3.20) define the two-dimensional critical manifold

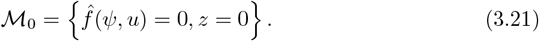

*ℳ*_0_ is normally hyperbolic if and only if 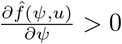 on *ℳ*_0_, where

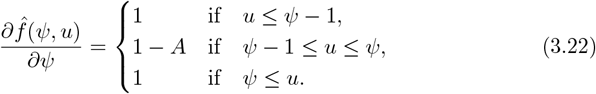

As the function 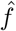 (3.6) piecewise smooth, we divide *ℳ*_0_ into three regions, according to the case distinction in the definition of 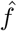. In the first region, where *ψ* ≤ *u*, we have

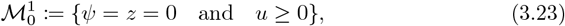

which is normally hyperbolic according to (3.22). For the second region, where *ψ*−1 ≤ *u ≤ ψ*, we find

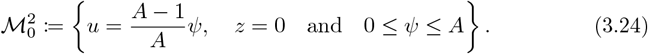

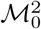 is normally hyperbolic for *A <* 1, but not normally hyperbolic for *A >* 1. Lastly, we consider the third region, where *u* ≤ *ψ* − 1. Here, the critical manifold

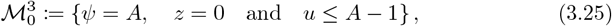

is again normally hyperbolic. In Figure 3.1, the critical manifold ℳ_0_ is shown projected onto the *ψu*-plane. The distinction between the case 0 *< A <* 1 and *A >* 1 can be observed. Based on the interpretation of *A* (3.7) as a measure of the interaction strength between morphogen and curvature dynamics, we will refer to 0 *< A <* 1 as the *weak interplay* regime, and to *A >* 1 as the *strong interplay* regime. In both regimes, we will study the existence of periodic patterns, see Sections 3.3 and 3.4.

In the fast system (3.8), we can again take the singular limit *ε* ↓ 0 to obtain

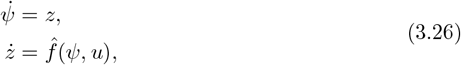

with *u* and *v* constant. The critical manifold ℳ_0_ (3.21) is the set of equilibria of the reduced fast system (3.26). We note that for 0 *< A <* 1, system (3.26) has a single saddle equilibrium. Conversely, for *A >* 1, there is a range of *u*-values for which (3.21) admits three equilibria; from the definition of 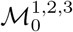, we see that these equilibria coexist when 0 *< u < A −* 1; see also Figure 3.1. The coexistence of hyperbolic equilibria in the reduced fast system (3.26) is a hallmark for nontrivial fast dynamics of periodic solutions in the full system (3.3). Therefore, we expect to find a clear qualitative distinction between periodic patterns in the weak (0 *< A <* 1) versus the strong (*A >* 1) morphogen-curvature interplay regime. This distinction will indeed be recovered in the analysis in Sections 3.3 and 3.4, and will also be clearly visible in the numerical simulations presented in Section 4.

**Figure 3.1:**
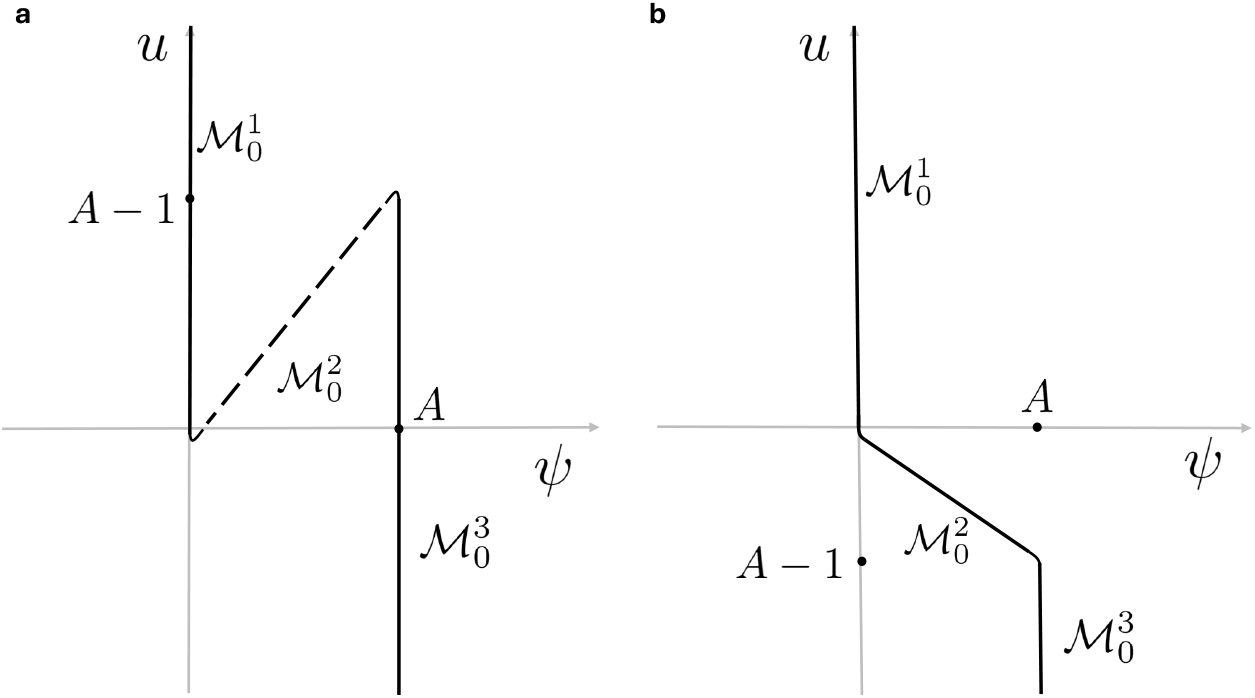
The projection of ℳ_0_ (3.21) onto (*ψ, u*)-space. Solid lines indicate normally hyperbolic regions, dashed lines indicate regions that are not normally hyperbolic. **a**. 0 *< A <* 1, here every value of *u*, the reduced fast system (3.26) admits only one equilibrium. **b**. *A >* 1, here three equilibria to (3.26) coexist for 0 *< u < A* − 1.

#### 3.2.1 Persistence of *ℳ*_0_

GSPT [20] provides the persistence of the normally hyperbolic regions of the critical manifold 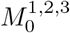 separately as invariant manifolds 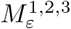 in the full system (3.5), provided 0 *< ε* ≪ 1 is sufficiently small. These invariant manifolds are *𝒪* (*ε*) close and diffeomorphic to their *ε* = 0 counterparts. In the case of (3.5), we note that *ℳss*_0_ (3.21) is itself invariant under the flow, which allows us to identify *ℳ*_*ε*_ = *ℳ*_0_. The piecewise smooth nature of 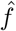 (3.6) restricts the direct application of GSPT to the separate smooth submanifolds 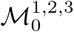. However, when we replace 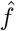 by a sufficiently smooth regularization as discussed in Section 2.3.1, the associated regularized, smooth manifold *ℳ*_0_ is fully normally hyperbolic when *A <* 1. Hence, it persists as an invariant manifold under the full flow of (3.5) for 0 *< ε* ≪ 1 sufficiently small.

However, when *A >* 1, the same regularization procedure causes 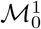 to lose normal hyperbolicity in an *𝒪* (*δ*_1_ + *δ*_2_) neighborhood of *u* = 0. Likewise, 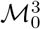 loses normal hyperbolicity in an 𝒪 (*δ*_1_ + *δ*_2_) neighborhood near its boundary at *u* = *A* −1. Hence, when *A >* 1, the choice of regularization – and in particular, the choice of the fixed regularization parameters *δ*_1_ and *δ*_2_ – restricts the region in phase space where persistent singular patterns can be constructed; see also Section 3.4.

### 3.3 Periodic patterns for weak morphogen-curvature interplay (*A <* 1)

In this subsection, we consider the weak morphogen-curvature interplay region – that is, we take 0 *< A <* 1. Our goal is to study the existence of stationary, spatially periodic patterns in system (1.4). For that reason, we look for periodic orbits in system (3.3), making use of the asymptotically small parameter 0 *< ε*≪ 1 that induces a scale separation in the dynamics of (3.3). Given such a periodic orbit, we then check whether it obeys geometric consistency condition (3.17).

#### 3.3.1 Existence of periodic orbits

The geometry of the critical manifold *ℳ*_0_ (3.21) is shown in Figure 3.1a. To study the dynamics on *ℳ*_0_, we consider the singular limit *ε* ↓ 0 of (3.5), to obtain

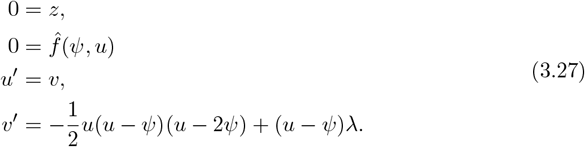

The first two equations restrict the flow to *ℳ*_0_ (3.21), while the last two equations determine the (slow) flow on ℳ_0_. Note that for 0 *< A <* 1, the projection of ℳ_0_ onto the (*ψ, u*)-plane can globally be written as a graph over *u*, see Figure 3.1a; hence, we can write *ψ* as a function of *u* by solving 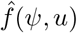 (3.6), yielding

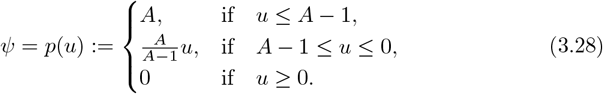

The slow dynamics on *ℳ*_0_ are therefore given by

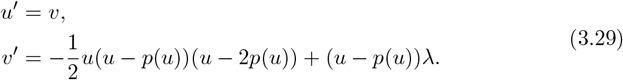

The equilibria of the piecewise smooth system (3.29) can be found by determining the intersections of the curve *ψ* = *p*(*u*) (representing the projection of *ℳ*_0_ onto the (*ψ, u*)-plane) with the nullclines *u* = *ψ* and 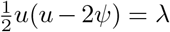. For *λ* ≤ 0, system (3.29) has a unique equilibrium at the origin, 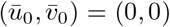. This equilibrium undergoes a pitchfork bifurcation at *λ* = 0, creating a pair of nontrivial equilibria (*ū*_*±*_, 0) with *ū*_+_ *≥* 0 and *ū* _−_ *<* 0. We find that

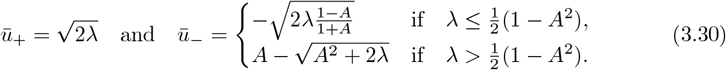

System (3.29) has Hamiltonian

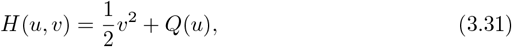

where

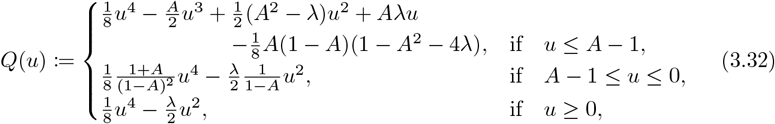

is gauged such that *Q*(*u*) is continuously differentiable. Using *H*(*u, v*) (3.31), we can determine the flow of (3.29). In Figure 3.2, the orbits of (3.29) are sketched for the two qualitatively different cases *λ <* 0 and *λ >* 0. We observe that in both cases, almost every orbit is periodic, the pair of homoclinics to the origin in the case *λ >* 0 being the only exception.

The geometry of ℳ_0_ and the slow dynamics on ℳ_0_ (see Figure 3.2) lead to the following result:

**Figure 3.2:**
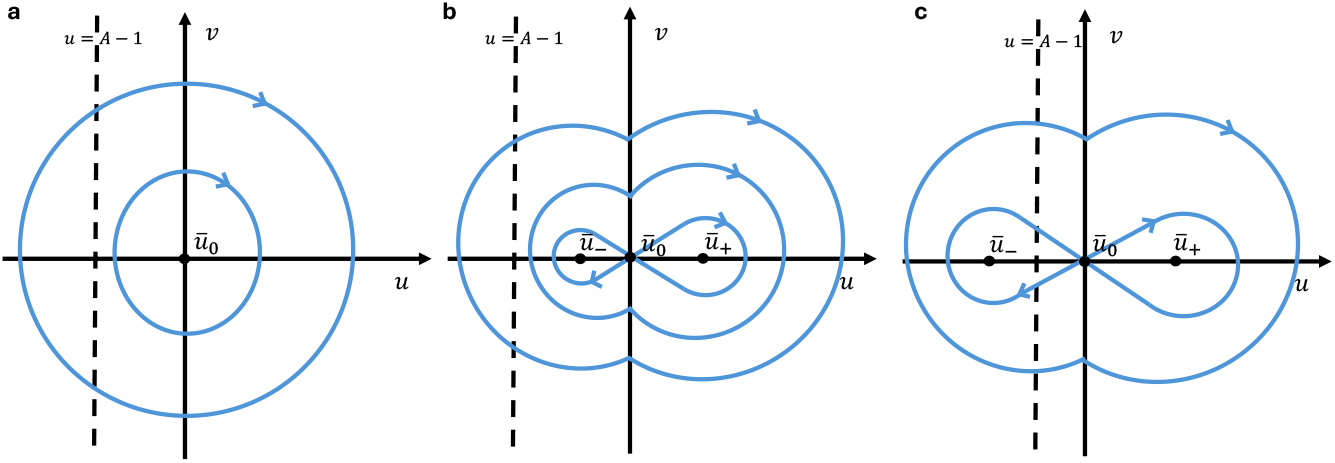
The slow flow (3.29) on *ℳ*_0_ (3.21) when 0 *< A <* 1, for different values of *λ*. **a**. *λ <* 0; **b**. 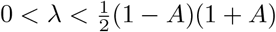**c**. 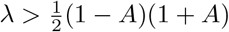.

##### Theorem 3.1.

*Let* 0 *< A <* 1, *and let* 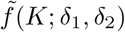 *be a smooth regularization of f* (*K*) (2.26). *For ε sufficiently small, system* (3.5) *exhibits a family, ℱ, of periodic orbits contained in ℳ*_0_ (3.21), *which are determined by the Hamiltonian* (3.31) *and every orbit in ℱ is a periodic orbit of the full system* (3.3). *Furthermore, there are no other periodic orbits in system* (3.3) *other than those contained in ℳ*_0_.

*Proof*. The critical manifold *ℳ*_0_ can be directly extended to its smoothly regularized counterpart by the same definition (3.21). Since for *δ*_1,2_ = 0 the separate segments 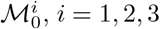 are normally hyperbolic, we can without loss of generality assume that the smooth regularization 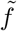 is such that the partial derivative 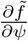 (3.22) is smooth and uniformly bounded away from zero, from which it directly follows that the smooth regularization of *ℳ*_0_ is fully normally hyperbolic. Hence, by standard Fenichel theory [22, Theorem 3.1.4], the manifold *ℳ*_0_ persists as an invariant, smooth, normally hyperbolic manifold *ℳ*_*ε*_ for sufficiently small *ε >* 0.

The dynamics on ℳ_0_ are given by (3.29). When *f* (*K*) is replaced by its smooth regularisation 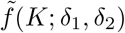, the piecewise smooth function *p*(*u*) (3.28) is mapped to its smooth regularised counterpart. Likewise, potential function *Q*(*u*) (3.32) in the Hamiltonian (3.31) is mapped to its smooth regularised counterpart. Hence, all piece-wise smooth periodic orbits to (3.29) are mapped to their smooth regularised counterparts. From the fact that the potential function *Q*(*u*) (3.32) behaves as *u*^4^ for large *u*, combined with the reversibility symmetry of system (3.29) and its Hamiltonian (3.31), we can conclude that system (3.29) admits a one-parameter family ℱ of periodic orbits that are parametrized by the value of the Hamiltonian on that orbit. By [6, Theorem 2.2], the reversibility symmetry of system (3.8) ensures that these periodic orbits persist for sufficiently small *ε >* 0.

Considering the reduced fast system (3.26), we note that for fixed *u*, system (3.26) has a single equilibrium of saddle type, which remains true when 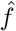 is replaced by its smooth regularization 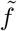 such that 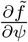 (3.22) is smooth and uniformly bounded away from zero. In addition, we observe that all non-equilibrium orbits of system (3.26) are unbounded. Since ℳ_0_ is normally hyperbolic and smooth, application of [22, Theorem 3.1.4] and [15, Theorem 8] implies that for *ε >* 0 sufficiently small, any orbit not contained in *ℳ*_0_ is unbounded and therefore cannot be periodic.

□

#### 3.3.2 Geometric consistency of the periodic orbits on ℳ_0_

In this section, we investigate the observability of the periodic orbits on ℳ_0_, drawn in Figure 3.2. As explained in Section 3.1, a periodic orbit in the steady state phase space of (3.5), might not be a solution to the PDE (1.4). Here we use geometric consistency condition (3.17),

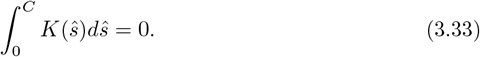

In this section, we assume that the Lagrange multiplier is negative (*λ <* 0), when *A <* 1, since the numerical simulation (see Section 4) indicates that this constraint is in order. The analysis for *λ >* 0 can be executed in a similar way. We first perform the consistency condition calculation for the piecewise smooth function *f* (*K*) (2.26) and for *ε* = 0 (see system (3.29)), then extend it for a smooth regularization 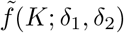 in Lemma 3.2.

Figure 3.2a shows the two possible types of periodic orbits for *λ <* 0 on *ℳ*_0_. In this parameter regime, there is one equilibrium, *ū*_0_ = 0, which is a center. Note that for any periodic orbit *H >* 0. We define *ū*_*p*_ (*ū*_*n*_) to be the unique positive (negative) value where the orbit intersects the positive (negative) u-axis; 2*H* − 2*Q*(*ū*_*p*_) = 0 (2*H*− 2*Q*(*ū*_*n*_) = 0). Furthermore, we define *Ĉ* as the period of the orbit. Since the “time” variable here is the curvilinear coordinate, *s*, there is a relationship between *Ĉ* and the arclength of the curve *C*, namely there exists a number *n* ∈ ℕ such that *C* = *n · Ĉ*. Now, without loss of generality, we assume that *u*(0) = *u*(*Ĉ*) = 0 and *v*(0) = *v*(*Ĉ*) *>* 0. Note that we can replace *C* by *Ĉ* in condition (3.33). In addition, we define *B* by the unique value where *u*(*B*) = 0 and 0 *< B < Ĉ*. By symmetry, 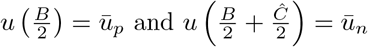.

To compute integral (3.33), we use the definition of *u* (3.2), *ψ* (3.2) and *K*_0_ (2.23), to derive *u* = *K* + *ψ*. Together with equation (3.28), we can now obtain the relationship between *u* and curvature *K*,

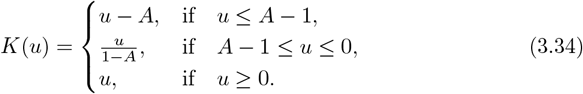

Note that these three different regions corresponding to the three sub-parts of the slow manifold, 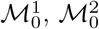 and 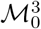. Using the definition of *Ĉ* and *B*, we can split the integral for *u ≥* 0 and *u* ≤ 0,

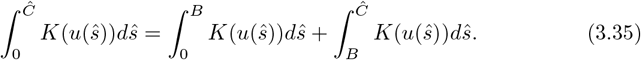

From the Hamiltonian (3.31) and the differential equation of *u*, (3.29), we derive

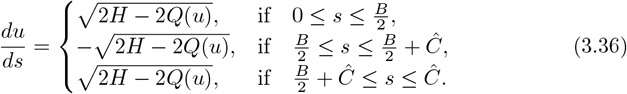

Firstly, we compute the first integral of equation (3.35) (for *u ≥* 0):

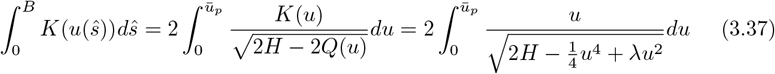

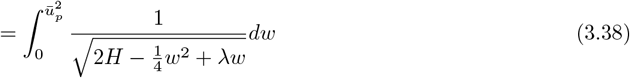

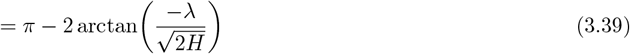

In (3.37) we use symmetry, 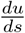 (3.36) and the definition of function *Q*(*u*) (3.32). In (3.38) we substitute *w* := *u*^2^. Lastly in (3.39), we use that

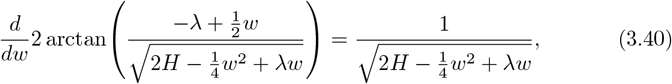

and

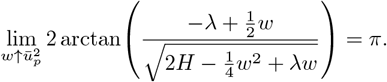

The computation of the second integral of (3.35) (for *u* ≤ 0) is more involved. Figure 3.2a shows that there are two different cases; orbits that are only on 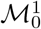 and 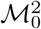 (when *A* − 1 ≤ *ū*_*n*_ ≤ 0) and orbits that partly lie on 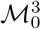(when *ū*_*n*_ ≤ *A* − 1).

We first investigate the first type of orbits, for which *A* − 1 *ū*_*n*_ 0. Note that this implies an upper bound for *H*. The integral over the curvature is computed as

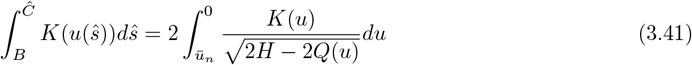

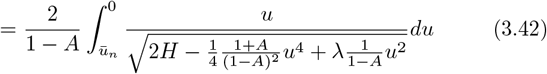

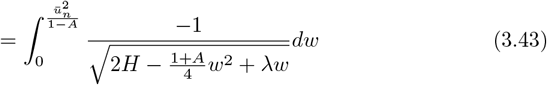

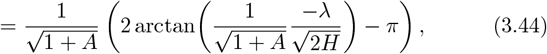

In (3.41) we use symmetry and 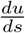 (3.36). In (3.42) the definition of function *Q*(*u*) (3.32) for *A* − 1 ≤ *u* ≤ 0 is substituted. To obtain (3.43) we substitute 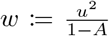.

Lastly (3.44) is attained using the antiderivative

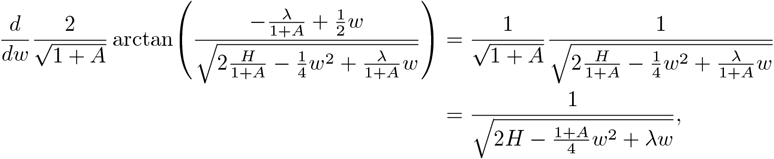

and

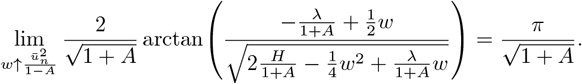

To investigate the second type of orbits, which partially lie on 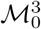, we take *ū*_*n*_ ≤ *A*−1.

The integral over the curvature is computed as follows:

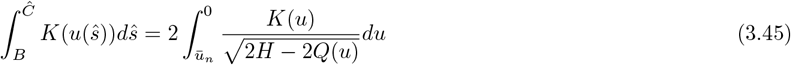

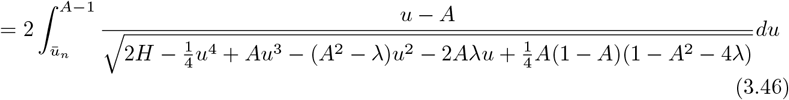

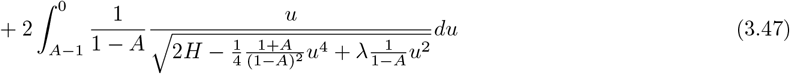

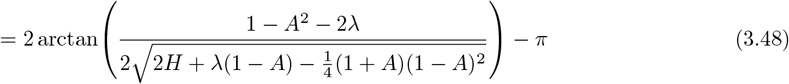

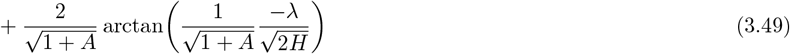

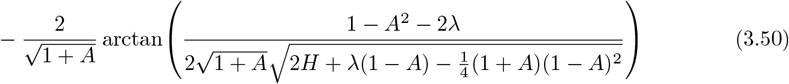

We use the above computations to prove Lemma 3.2.

##### Lemma 3.2.

*Let* 0 *< A <* 1, *λ <* 0 *and consider a periodic orbit* (*u*_*p*_, *v*_*p*_) *in system* (3.29). *Let* 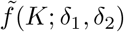 *be a smooth regularization of f* (*K*) (2.26) *with δ*_1,2_ *sufficiently small. For ε sufficiently small, the perturbed counterpart* 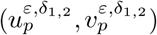*does not meet geometric consistency condition* (3.17) *and is therefore not observable as a solution of the PDE* (1.4).

*Proof*. A periodic orbit (*u*_*p*_, *v*_*p*_) in system (3.29) intersects the *u*-axis at *ū*_*n*_ and at *ū*_*p*_, with *ū*_*n*_ *< ū*_*p*_. We are interested in computing the integral over the curvature for the perturbed counterpart 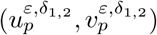. By Theorem 3.1, every periodic orbit of system (3.5) is contained in the invariant manifold *ℳ*_*ε*_. Therefore, every periodic orbit of system (3.5) can to leading order in *δ*_1,2_ and *ε* be described as a periodic orbit of system (3.29), which gives the leading order (in *δ*_1,2_) description of the dynamics on the smoothly regularized *ℳ*_0_.

We first consider the case *A*−1 ≤ *ū*_*n*_ < 0. By rewriting (3.39) and (3.44), we can deduce

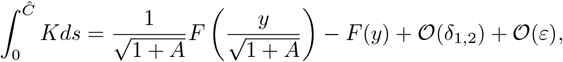

for *F* (*x*) := 2 arctan(*x*) − *π* and 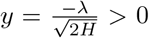, since the periodic orbit we are integrating over is to leading order equal to its *δ*_1,2_ = *ε* = 0 counterpart. We can derive the following properties:

- For 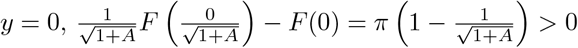,
- 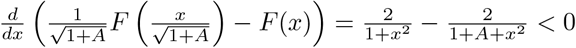,
- 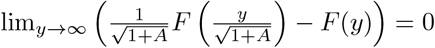,

to conclude that 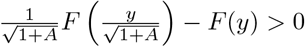 for all *y ≥* 0.

Now, for fixed *λ* and *H*, we can take *δ*_1,2_ and *ε* sufficiently small such that 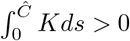.

Next, we consider the case *ū*_*n*_ *< A−* 1. The integral over the curvature for *û*_*n*_ *< A −*1 equals the sum of (3.39) and (3.48)-(3.50) to leading order. By rewriting these equations, we identify 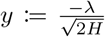 and 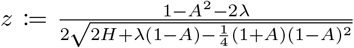 and function 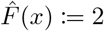arctan(*x*) such that

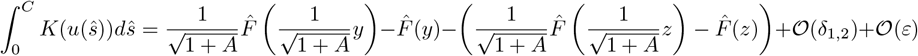

since again the periodic orbit we are integrating over is to leading order equal to its *δ*_1,2_ = *ε* = 0 counterpart. For 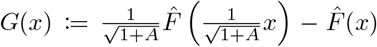, we identify the following properties:

- *G*(0) = 0,
- 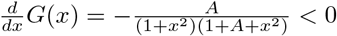
- 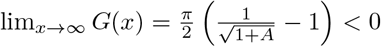

We deduce that *G*(*y*)−*G*(*z*) = 0 implies *y* = *z*, resulting in 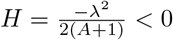. However, since *H >* 0 for the orbits we are investigating, *y* ≠ *z* in the feasible regime. For *y* ≠ *z*, we can take *δ*_1,2_ and *ε* small enough such that *G*(*y*) − *G*(*z*) + (*δ*_1,2_) + 𝒪(*ε*) ≠ 0.

Since (3.17) was formulated in Section 3.1 as a necessary condition for a solution to be observable, we conclude that when *δ*_1,2_ *>* 0 are sufficiently small, and when *ε >* 0 is sufficiently small, the periodic counterpart 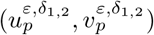 of (*u, v*) is not observable as a solution of the PDE (1.4). □

**Figure 3.3:**
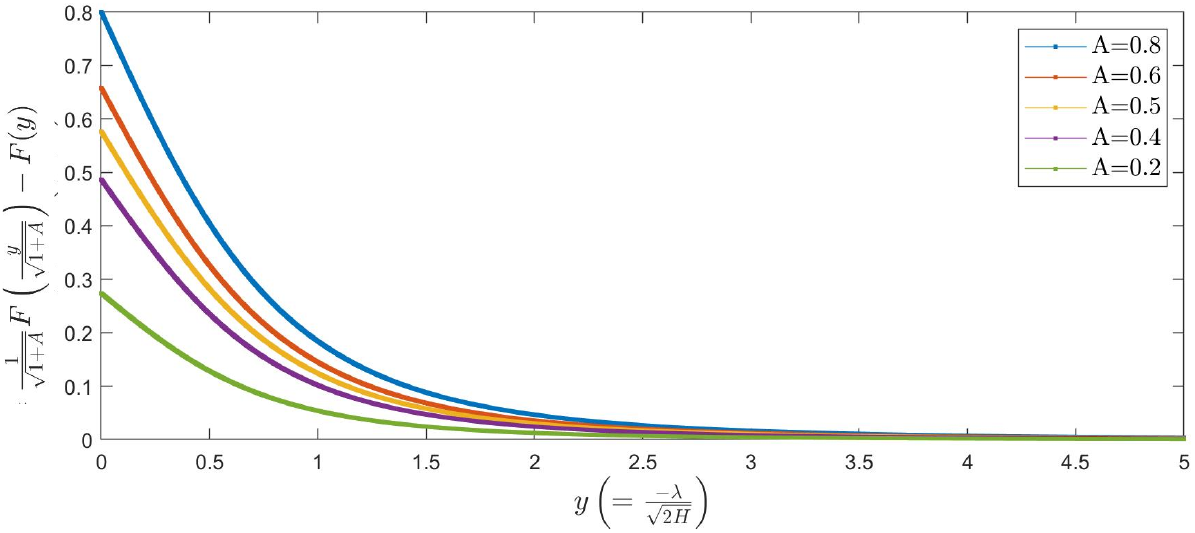
The *𝒪* (1) terms of the integral of the curvature, 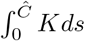, of a periodic orbit on *ℳ*_*ε*_ for 0 *< A <* 1 and *A* − 1 ≤ *ū*_*n*_ *<* 0, displayed as a function over 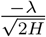. See proof of Lemma 3.2 for the properties of this function.

See Figure 3.3 for a plot of 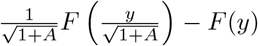, for some values of *A*. Note that the integral value rapidly decreases when − *λ* is large or when *H* is small.

We can interpret the results of Lemma 3.2 as follows. For an orbit (which describes the curvature of a curve over the curvilinear coordinate) to be observable, it needs to have “as much negative curvature as positive curvature”. The part of the orbit on 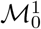 describes the curve when no morphogen is present. Here, the curve has positive curvature (is curving outward). The part of the orbit on 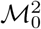 (and 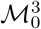) describes the curve when there is morphogen present. Here, the curvature is negative. Our analysis now shows that the dynamics on and the geometry of the slow manifold make it impossible for the total amount of negative curvature to be sufficient to balance the total amount of positive curvature collected on 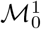. Therefore, it is impossible to start and end with the same slope of the curve (i.e. to obey the periodic boundary conditions (2.28)–(2.30)). Still, system (1.4) might still produce steady state periodic solutions as described in 3.1, when *ε* and *δ*_1,2_ are small but not too small, see Section 3.5.

### 3.4 Periodic orbits for strong morphogen-curvature interplay (*A >* 1)

In this subsection, we consider the strong morphogen-curvature interplay region, which is characterized by *A >* 1. As in the previous subsection, our goal is to study the existence of stationary, spatially periodic patterns in system (1.4), by identifying periodic orbits in system (3.3). Again, the asymptotically small parameter 0 *< ε* ≪ 1, which induces a scale separation in the dynamics of (3.3), will play a pivotal role. For the periodic orbits that we find, we check the geometric consistency conditions in Section 3.1 to validate the observability of these orbits. For most of the computations, we will consider the piecewise smooth function *f* (*K*) (see Section 3.4.2 and 3.4.3). The results, however, can be analogously converted to system with a the smooth regularization of *f* (*K*), namely 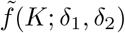.

#### 3.4.1 Geometry of the critical manifold

The geometry of the critical manifold *ℳ*_0_ (3.21) is shown in Figure 3.1b. In contrast to the weak interplay regime, 0 *< A <* 1, the manifold *ℳ*_0_ intersects the hyperplane {*u* = *u*_0_} more than once, yielding multiple equilibria in the reduced fast system (3.26). We will show that this allows for the existence of bounded orbits that ‘jump’ between different sheets of _0_. This situation is fundamentally different from the case 0 *< A <* 1, where all bounded orbits are fully contained in the slow manifold; see Section 3.3. To study and construct periodic orbits for *A >* 1, we consider the reduced slow and fast dynamics in more detail.

#### 3.4.2 Reduced fast dynamics

Using the definition of 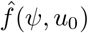 (3.6), we restate (3.26) for convenience:

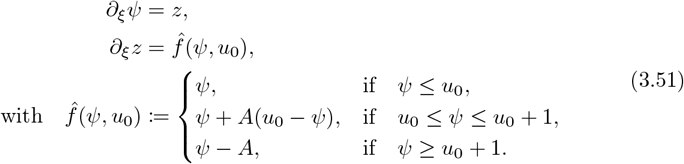

System (3.51) is a combination of three linear dynamical systems in three separate subsystems, with a continuous transition at the values *ψ* = *u*_0_ and *ψ* = *u*_0_ + 1. The system is Hamiltonian, with the piecewise smooth Hamiltonian function

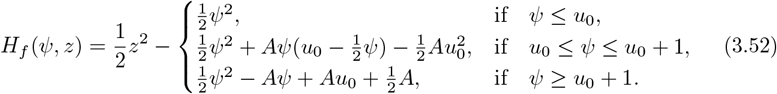

**Figure 3.4:**
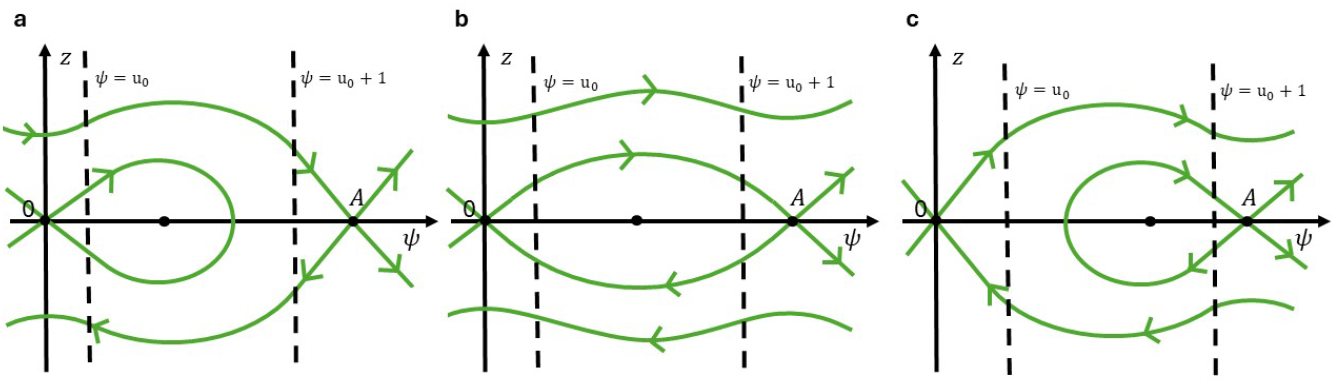
The phase plane of system (3.51) for three representative values of *u*_0_. **a**. For 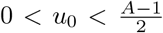, there is a homoclinic orbit to the origin (0, 0). 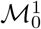 **b**. For 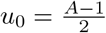, a pair of heteroclinic orbits between the saddle equilibria (0, 0) and (*A*, 0) exists. **c**. For 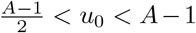, there is a homoclinic orbit to the saddle equilibrium (*A*, 0).

*H*_*f*_ is gauged such that *H*_*f*_ (0, 0) = 0, and such that it is continuously differentiable in *ψ* at the transition values *ψ* = *u*_0_ and *ψ* = *u*_0_ + 1.

The number of equilibria of system (3.51) depends on the value of *u*_0_. Solving 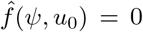 under the assumption that *A >* 1, we find that for *u*_0_ *<* 0, there is a single saddle equilibrium at (*ψ, z*) = (*A*, 0); for *u*_0_ *> A* − 1, there is a single saddle equilibrium at the origin. When 0 < *u*_0_ *< A*−1, system (3.51) admits three equilibria: a saddle at the origin, a saddle at (*ψ, z*) = (*A*, 0), and a center at 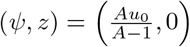 see also Figure 3.1b.

The structure of the stable and unstable manifolds 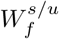 of the saddles (0, 0) and (*A*, 0) is determined by the level sets of the Hamiltonian *H*_*f*_. We consider *u*_0_-values for which three equilibria coexist, i.e. 0 *< u*_0_ *< A* − 1. We find

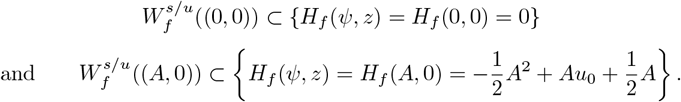

It follows that 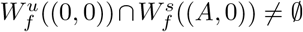 if and only if 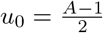 ; symmetry implies that also 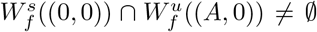 for that choice of *u*_0_. In other words, there exists a pair of heteroclinic orbits in system (3.51) that connects the two saddles (0, 0) and (*A*, 0) if and only if 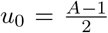. By continuity, the structure of the level sets of *H*_*f*_ for other values of 0 *< u*_0_ *< A* − 1 follows. A sketch of the phase plane of (3.51) for representative values of *u*_0_ is given in Figure 3.4. As the critical manifold 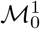 (3.23) is given by *{ψ* = *z* = 0, *u ≥* 0*}*, its stable and unstable manifolds are given as a union over *u ≥* 0 and *v* ∈ ℝ of the stable and unstable manifolds of the origin in the fast reduced system (3.51), i.e. 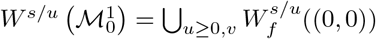. Similarly the stable and unstable manifolds of the normally hyperbolic branch 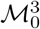 (3.25) are given by 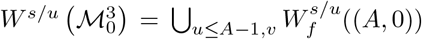. We know by the above arguments that 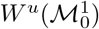 and 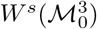 intersect, and that this intersection is contained in the hyperplane 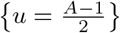 ; the same holds for the intersection of 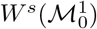 and 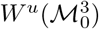. The structure of the level sets of *H*_*f*_ (3.52) reveals that both intersections are transversal, see also Figure 3.5.

**Figure 3.5:**
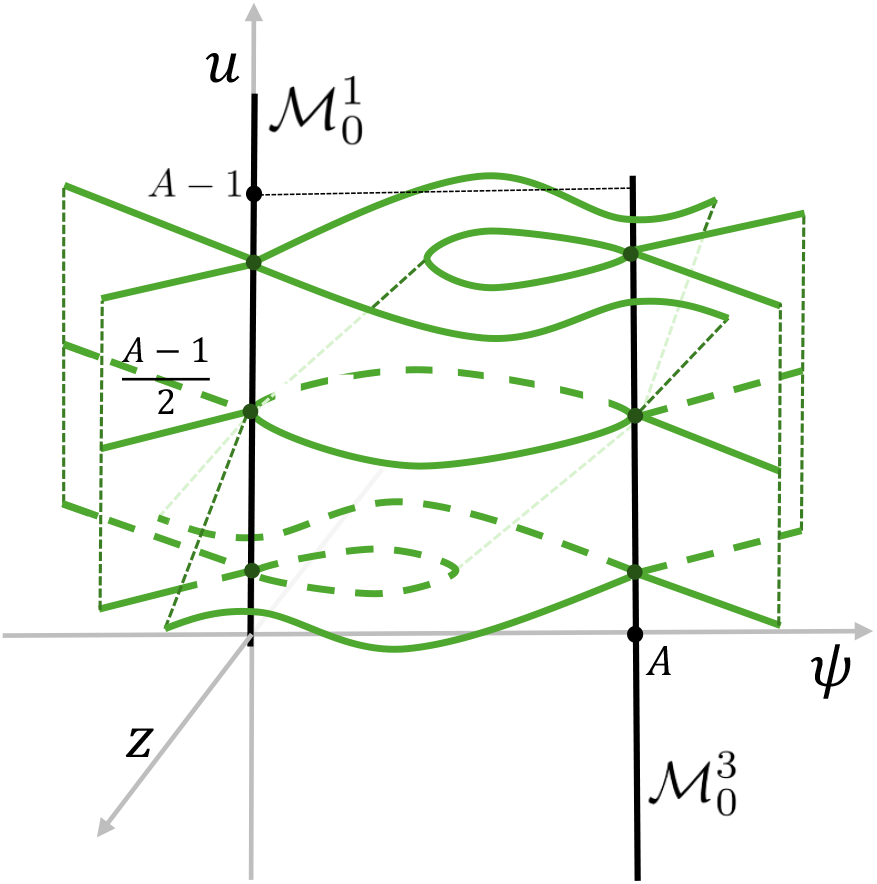
Sketch of the stable and unstable manifolds of 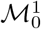 and 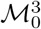 for the fast dynamics (3.51) for varying *u* = *u*_0_.

#### 3.4.3 Slow reduced dynamics

We determine the flow on the critical manifold *ℳ*_0_ (3.21). Since 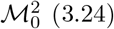 (3.24) is not normally hyperbolic, and therefore does not (necessarily) persist for 0 *< ε* ≪ 1, we restrict our analysis to the normally hyperbolic branches 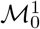 (3.23) and 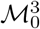 (3.25).

The reduced slow dynamics on 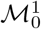, where *ψ* = *z* = 0, are given by

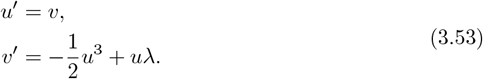

The number of equilibria of system (3.53) is determined by the sign of *λ*. For *λ <* 0, (3.53) has a single equilibrium at the origin (*u, v*) = (0, 0) of center type. For *λ >* 0, (3.53) has three equilibria: a pair of centers at 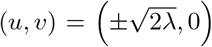 and a saddle at the origin. Note that the dynamics on 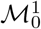 are, by its definition (3.23), restricted to the half-plane *u ≥* 0. System (3.53) is Hamiltonian, with Hamiltonian function

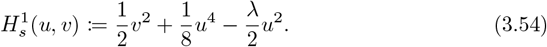

The level sets of 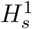 can be used to determine the orbits of (3.53); see also Figure 3.6a. For future reference, we denote the one-dimensional stable and unstable manifolds of (0, 0) of the reduced slow flow (3.53) by 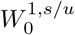 ((0,0)). These manifolds are nonempty when *λ >* 0, see again Figure 3.6a. Furthermore, we note that equilibria of the reduced slow system (3.53) correspond to equilibria of the full four-dimensional system (3.5). Hence, for future reference, we define

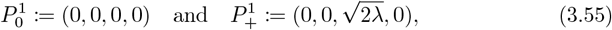

and indicate their position in Figure 3.6a.

The reduced slow dynamics on 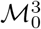, where *z* = 0 and *ψ* = *A*, are given by

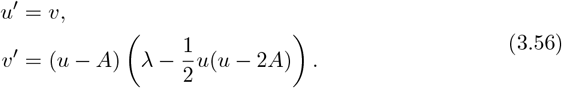

The dynamics of system (3.56) are similar to the dynamics on 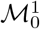 (3.53); indeed, system (3.53) can be transformed into system (3.56) under the substitution *u* → *u* − *A*, 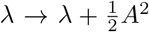. Hence, for 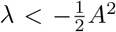, system (3.56) has a single center equilibrium at (*u, v*) = (*A*, 0). For 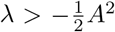, system (3.56) has three equilibria: one saddle at (*A*, 0) and a pair of centers at 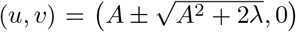. Note that the dynamics on 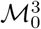 are, by its definition (3.25), restricted to the half-plane *u* ≤ *A* − 1; therefore, the equilibria (*A*, 0) and 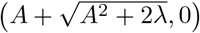 are not contained in 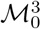. The center equilibrium 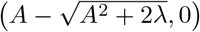 is contained in 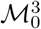 if and only if 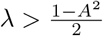.

System (3.56) is Hamiltonian, with Hamiltonian function

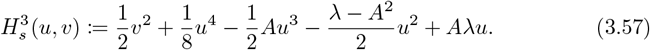

The level sets of 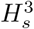 can be used to determine the orbits of (3.56); see also Figure 3.6b. For future reference, we define the equilibrium of the full system (3.5)

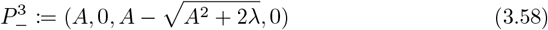

and indicate its position in Figure 3.6b.

**Figure 3.6:**
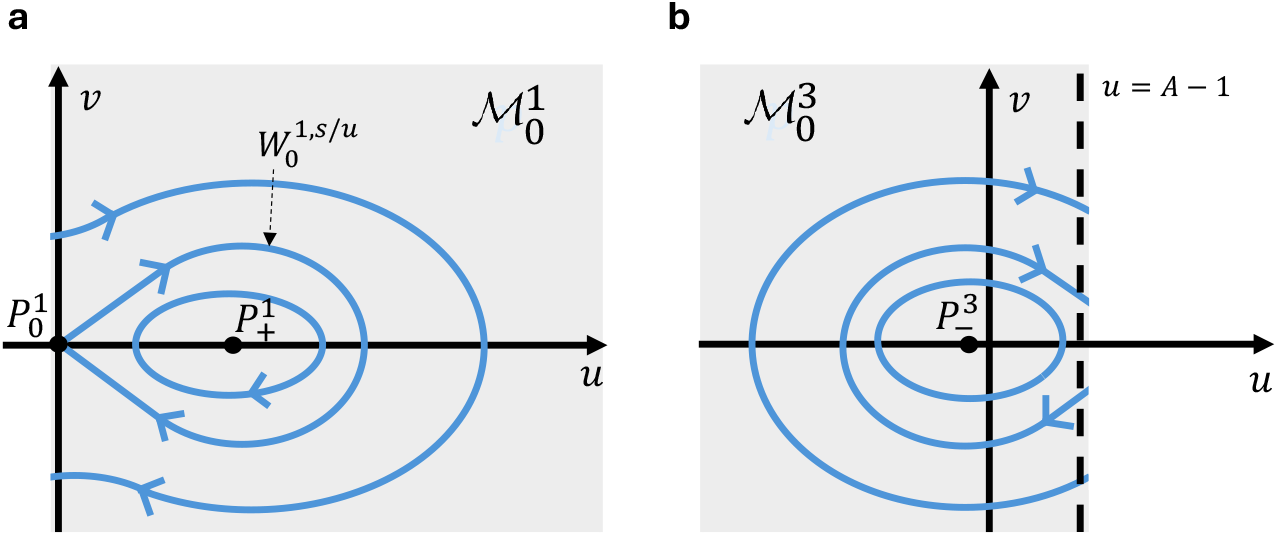
Sketch of the slow dynamics (3.53)/(3.56) on the critical manifolds 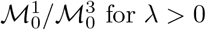.

#### 3.4.4 Existence of periodic orbits: setup

To study the existence of periodic orbits in system (3.5) for *A >* 1, we follow a similar approach to the case 0 *< A <* 1, see Section 3.3. In contrast to the weak interaction case, we now have the opportunity to construct three qualitatively different types of periodic orbits (see Figure 3.5): those that are fully contained in either branch 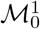 or 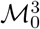of the critical manifold, those that partly lie close to either 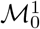 or 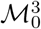, but exhibit ‘fast homoclinic jumps’, or those that exhibit ‘fast heteroclinic jumps’ between these branches and thus have part of the orbit lie close to 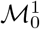, and part close to 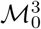. It should be noted that as long as 0 ≤ *u* ≤ *A* − 1, the location and geometry of the slow manifolds 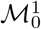 and 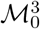 do not change qualitatively when we take smooth regularization 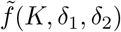. Furthermore, note that the curvature of any periodic orbit that is fully contained in 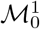, is strictly positive *K* = *u >* 0. Moreover, any periodic orbit that is fully contained in 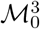, is strictly negative *K* = *u* −*A < A* −1 −*A <* 0. Hence, these orbits cannot comply with geometric consistency condition (3.17) and therefore can never be observable as solutions to the PDE (1.4b). Furthermore, other possible orbits in this system, like purely fast periodic orbits or periodic orbits with a homoclinic jump in the fast system do not show up in any of the numerical simulation see Section 4. In this paper, we therefore restrict our analysis to the ‘heteroclinic slow-fast’ type of periodic orbits when *A >* 1.

The geometry of the reduced slow flows (3.53) and (3.56) is determined by the value of *λ*, see section 3.4.3. Depending on the value of *λ*, several concatenation configurations are possible. In this paper, we focus our attention on the case *λ >* 0, and prove the existence of a slow-fast orbits in the following subsections. For other ranges of *λ*, the existence of periodic orbits can be proven using similar means, see e.g. [18; 7]. Our choice for *λ >* 0 is motivated by the numerical simulations of the full PDE system (1.4), see Section 4.

Before we prove the existence of these periodic orbits, we first prove the existence of an orbit, *γ*_*h*_(*s*), homoclinic to the steady state 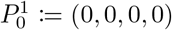 “jumping” to 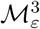 and back, since it helps in setting up the more elaborate proof of the existence of periodic orbits, see Theorem 3.4. To prove the existence of the homoclinic and period orbits, we utilize techniques from GSPT like singular skeleton and take off and touch down curves, see [15; 20; 22; 18] for more background information and theory. Note that in the proof for homoclinic orbits we explicitly use the smooth regularization, 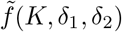, since 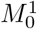 is not normally hyperbolic in an *ϵ* region around 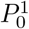 when we use piecewise smooth function *f* (*K*).

#### 3.4.5 Existence of homoclinic orbits to 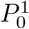, jumping to 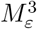 and back

To prove the existence of ‘slow-fast’ homoclinic orbit, we construct the so-called singular skeleton *𝒮*_*h*_ using the reduced slow dynamics on 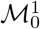and 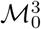, combined with the reduced fast dynamics that govern the flow in between 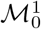 and 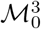. However, to show persistence of 𝒮_*h*_ for *ε >* 0, some extra work is required. Namely, the existence of a heteroclinic connection in the fast dynamics, system (3.51), does not automatically guarantee a non-empty intersection of 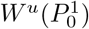 and 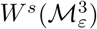 for *ε >* 0. Furthermore, *γ*_*h*_ touches down on, but also takes off from 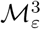. Therefore, *γ*_*h*_ does not lie in 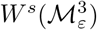. We use the persistence of the intersection 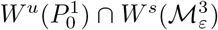 and the reversibility of the system to prove that there is an orbit exponentially close that has the properties we desire. Below, we go over the technical details and prove in Theorem 3.3 the existence of a homoclinic orbit for *ε >* 0 that is *𝒪* (*ε*) close to *𝒮*_*h*_.

**Figure 3.7:**
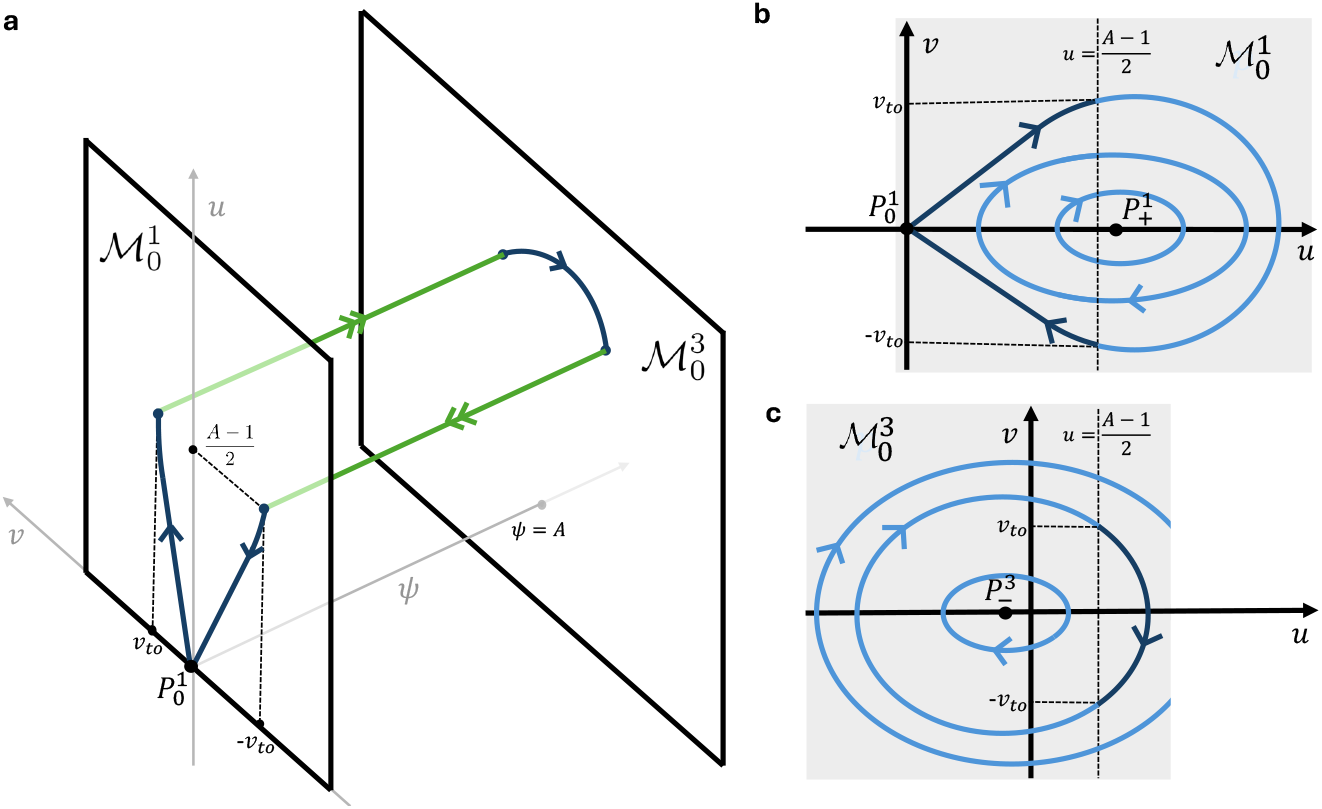
The concatenation of slow and fast orbit segments, to form a homoclinic orbit to *P*_0_, for 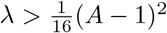. The jumping points are indicated by *±v*_*to*_ and 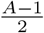. **a**. Sketch of the singular skeleton projected on *ψuv*-space. **b/c**. Slow dynamics on 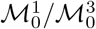, see system (3.53)/(3.56). Dark blue indicates the part of the orbit selected for the singular skeleton.

##### Singular skeleton *𝒮*_*h*_ **of homoclinic orbit** *γ*_*h*_

The singular skeleton 𝒮_*h*_ consists of five sub-parts. For each subpart we either “freeze” the fast dynamics and follow the slow flow on the critical manifold or we “freeze” the slow dynamics and follow the fast dynamics. See Figure 3.7 for a graphical representation of the singular skeleton in *ψuv*-space. In Section 3.4.2 and 3.4.3 we analyze the fast and slow reduced systems with the piecewise smooth function *f* (*K*) that provides an 𝒪 (1) approximation of the flow for the system with smoothly regularized function 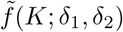.

For the first part of the singular skeleton *𝒮*_*h*_, we follow the slow reduced flow on 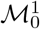 via 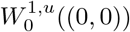 until (*u, v*) = (*u*_*to*_, *v*_*to*_). Recall that 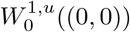 is the unstable manifold of 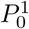 for the slow reduced dynamics. The second part of *𝒮*_*h*_ follows the fast dynamics to jump from 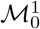 to 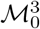. This orbit lays in 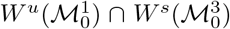, see Figure 3.5. Using the Hamiltonian of the fast dynamics, *H*_*f*_ (*ψ, z*) (3.52), we compute that for 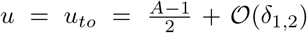, such a heteroclinic connection exists. Note that we can use the Hamiltonian 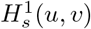 in (3.57), to compute that 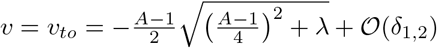. On 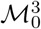, we follow the slow flow, (3.56), on level curve 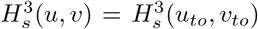until (*u, v*) = (*u*_*to*_, −*v*_*to*_). Lastly, for the fourth and fifth part of *𝒮*_*h*_ we jump from 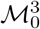 back to 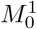 via the heteroclinic fast connection and use 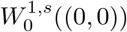 to finish this homoclic singular skeleton. Note that from the computation of *u*_*to*_, we can derive an extra bound on parameter *λ*. Namely, from Hamiltonian (3.54), it follows that 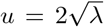 to leading order when the homoclinic orbit of 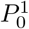 in the reduced slow dynamics (3.53) intersect the *v* = 0 axis. To be able to construct a fast jump, we need that 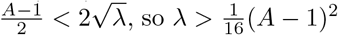.

##### Persistence of intersection of approach of 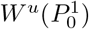 and 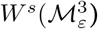 for *ε >* 0

In Section 3.2, we analyzed the location and the type of the critical manifolds. Recall that by Fenichels first theorem there exist invariant manifolds 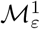 and 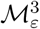 that are *𝒪* (*ε*) close and diffeomorphic to 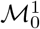 and 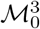 respectively. By Fenichels second theorem, we find that there exist stable and unstable manifolds 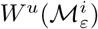 and 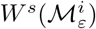 that are *𝒪* (*ε*) close and diffeomorphic to 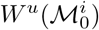 and 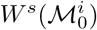 respectively, for *i* ∈ *{*1, 3}. In Section 3.4.2, we analyze the fast reduced system. These results can be transferred to the system with smooth regularization 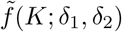, showing that 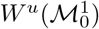 and 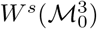 as well as 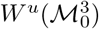 and 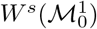 intersect transversely, when 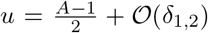. Therefore, we can conclude that for *ε* small enough, this intersection persists. Note that 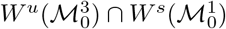) is two-dimensional.

Since 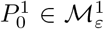, we have that 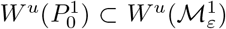. The intersection 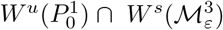 is a one-dimensional subset of 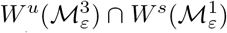. This intersection exactly describes an orbit, 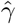, that goes to 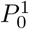 in backward time and touches down on 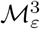 in forward time, i.e., it does not depart from 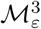 again.

##### The touch down curve of 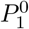 on 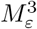

For *ε >* 0, the homoclinic orbit *γ*_*h*_, follows at leading order in *ε* the flow 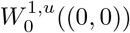. It “takes off” from 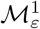 at (*u, v*) = (*u*_*to*_, *v*_*to*_) and follows the fast flow described by (3.51). At leading order, the slow varying variables, *u* and *v* do not change during this fast jump. Therefore, the orbit “touches down” on 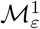 at (*u, v*) = (*u*_*to*_, *v*_*to*_) at leading order. As a function of parameter *λ*, the touchdown curve *𝒯*_*d*_(*λ*) is given by

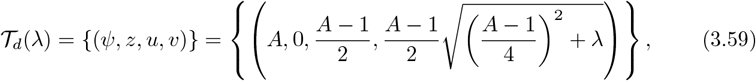

to leading order (in *ε, δ*_1,2_).

##### Symmetry of the full system

The full system of ODEs, system (3.5)/(3.8) accommodates a reflectional symmetry. If we send

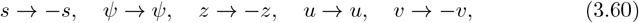

the system of equations does not change. Meaning, any orbit in the four dimensional phase space must have a symmetric counter (piece of) orbit. Moreover, when an orbit crosses exactly through the hyperplanes *v* = 0 and *z* = 0 at the same “time” (i.e. through {*v* = *z* = 0}), the orbit must be symmetric itself. We use this property of the system in the proof of the existence of a homoclinic orbit. Not that the slow and fast dynamics also exhibit (a reduced version of) this symmetry in two coordinates.

###### Theorem 3.3.

*Fix A >* 1, 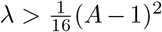 *and let* 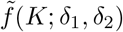 *be a smooth regularization of f* (*K*) (2.26). *For ε >* 0 *sufficiently small, system* (3.5) *exhibits a homoclinic orbit, γ*_*h*_(*ξ*) = (*ψ*_*h*_(*ξ*), *z*_*h*_(*ξ*), *u*_*h*_(*ξ*), *v*_*h*_(*ξ*)) *to* 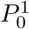 *that merges with the skeleton 𝒮*_*h*_ *as ε* ↓ 0.

For this proof, we follow the approach of Jaibi et al. [18] for multi-front patterns that was based on the geometric set-up of [7] for multi-pulse patterns. The general idea is to track orbits *γ*(*ξ, d*) that are in 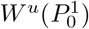, and very close to the orbit 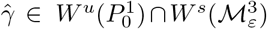. These orbits follow the dynamic behavior of 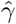, however they are not asymptotic to 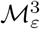 as *ξ*→ ∞. By selecting an orbit that passes through *v* = 0 and *z* = 0 at the same moment in “time” (i.e. *ξ*) we can use the symmetry of the system to conclude that the homoclinic orbit exists.

*Proof*. Let 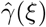 be the orbit in the intersection 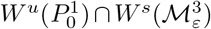) for *ε >* 0, that we have shown to persist. In leading order of *ε*, 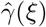 is described by the fast dynamics for 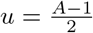, therefore 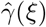 intersects 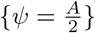 transversally. We name this intersection point *Q*. Note that 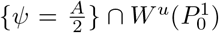 describes a curve, *𝒞*, where *Q* ∈ *𝒞*. Curve *𝒞* provides a parametrization of orbits 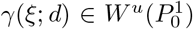 that are “close enough” to 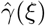, where *d*_−_ *< d < d*_+_ is the arclength coordinate along *𝒞*, such that 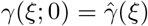 The saddle structure around 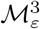 divides {*γ*(*ξ*; *d*), *d*_−_ *< d < d*_+_} in two. Without loss of generality we choose that for *d*_−_ *< d <* 0, *γ*(*ξ*; *d*) leaves the neighborhood of 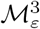 for the same sign of *z, z <* 0. For 0 *< d < d*_+_, *γ*(*ξ*; *d*) passes through *z* = 0 and leaves the neighborhood of 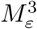 for *z >* 0. Here we used that 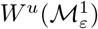 is *𝒪* (*ε*) close and diffeomorphic to 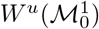, see Figure 3.4b.

Thus, we have a one-parameter family of orbits {*γ*(*ξ*; *d*), 0 *< d < d*_+_} that stay close to 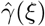 for some “time” *ξ* (by continuous dependence on initial conditions). For *d* sufficiently small, orbits *γ*(*ξ*; *d*) follow the fast flow, touch down on 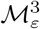 asymptotically close to the touch down point *𝒯*_*d*_(*λ*) and subsequently follow the slow flow on 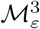, at leading order determined by the level curve 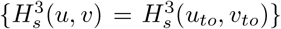 of (3.56). Therefore, for *d* small enough, *γ*(*ξ*; *d*) passes through {*v* = 0} for some *ξ*^∗^ = *ξ*^∗^(*d*), with *γ*(*ξ*^∗^(*d*); *d*) exponentially close to 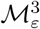.

By decreasing *d*, i.e. by bringing *γ*(*ξ*; *d*) closer to 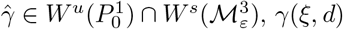 will remain (exponentially) close to 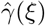, and thus to 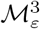, for an increasing period of “time” *ξ*. Clearly, 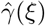 does not depart from 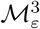 and thus never crosses through {*z* = 0}. However, for *d >* 0 any orbit *γ*(*ξ*; *d*) will eventually take off from 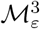. Thus, any orbit *γ*(*ξ*; *d*) will also eventually cross through {*z* = 0}. By taking *d* sufficiently close to 0 the moment *ξ*^*†*^(*d*) this intersection takes place can be postponed until after *γ*(*ξ*; *d*) has passed through {*v* = 0}, i.e. *ξ*^*†*^(*d*) *> ξ*^∗^(*d*) for *d* sufficiently small.

On the other hand, we may also take *d* relatively large, so that *γ*(*ξ*; *d*) only remains exponentially close to 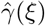 for a relatively short amount of time and takes off from 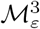 before 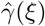 crosses through {*v* = 0}. Before the moment of taking off, *γ*(*ξ*; *d*) must have crossed through {*z* = 0} which implies that *ξ*^*†*^(*d*) *< ξ*^∗^(*d*) for these values of *d*. It follows by continuity that there must be a *d*^∗^ such that *ξ*^*†*^(*d*^∗^) = *ξ*^∗^(*d*^∗^) which implies that *γ*(*ξ*; *d*^∗^) passes through {*z* = 0} and {*v* = 0} at the same moment in time. By the symmetry of the system, (3.59), we now have found the orbit *γ*_*h*_(*ξ*) = *γ*(*ξ*; *d*^∗^).

□

#### 3.4.6 Existence of periodic orbits jumping between 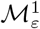 and 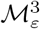

For the existence of periodic orbits we can use a very similar approach: first we construct a singular skeleton, then we establish the existence of a periodic orbit that merges with this skeleton in the limit *ε* → 0 by following the geometric set-up of [7].

##### Singular skeletons 𝒮_*p*_(*H*_*f*_)

Figure 3.8 shows a graphical representation of a singular skeleton of a periodic orbit that jumps between 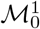 and 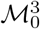. We can find a one-parameter family of singular skeletons 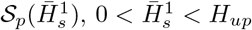, where 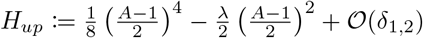, the Hamiltonian value for which the orbit crosses the line 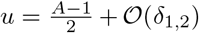 at the same time as the *u*-axis. Namely, setting the Hamiltonian of the slow reduced flow on 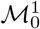 (3.54) equal to 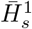 uniquely fixes an orbit piece on 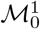. Denote *v*_*j*_ and −*v*_*j*_ by the values of *v* where this orbit piece intersects the line 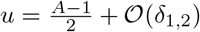. The singular skeleton consists of a part on 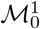, described by 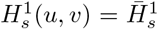 (3.54) and dynamics (3.53). Then, when *v* = *v*_*j*_, it jumps to 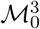, using the fast reduced dynamics (3.51) for the regularized function 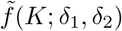, keeping *u* and *v* constant. On 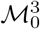, we follow an orbit part described by the slow reduced dynamics (3.56). When *v* = −*v*_*j*_, we jump back to 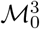 via the fast dynamics, completing the singular skeleton of the periodic orbit. Note that from the geometric construction *v*_*j*_ *< v*_*to*_. Furthermore, we again need that 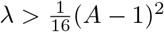.

###### Theorem 3.4.

*Let* 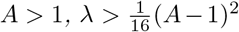, *let* 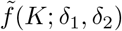 *be a smooth regularization of f* (*K*) (2.26) *and fix* 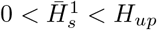. *For ε >* 0 *sufficiently small, system* (3.5) *exhibits a one-parameter family of periodic orbits*, 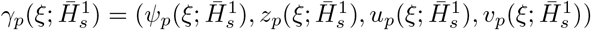 *that merge with skeleton* 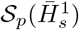 *as ε* ↓ 0.

**Figure 3.8:**
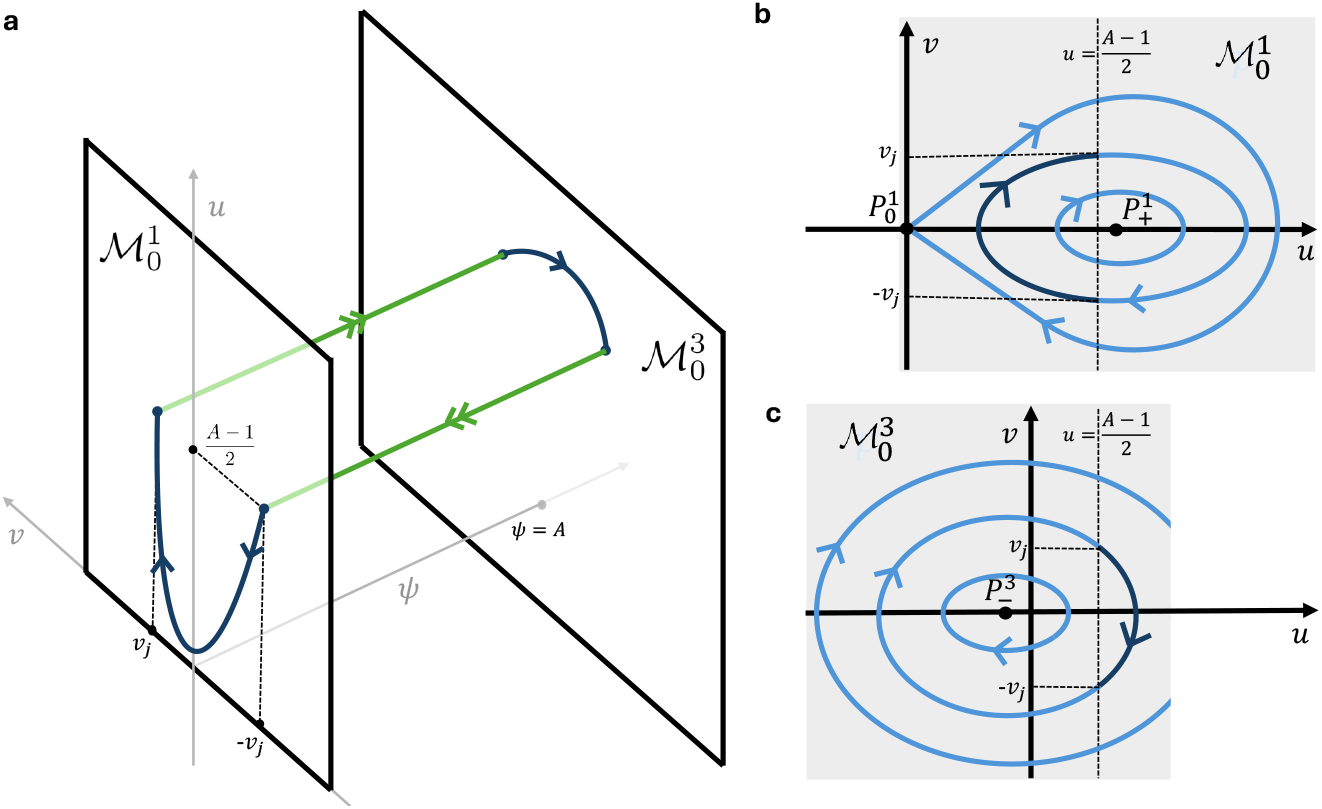
Example of the concatenation of slow and fast orbit segments, to form a periodic orbit, for 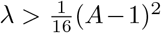. The jumping points are indicated by *±v*_*j*_ and 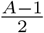. **a**. Sketch of the singular skeleton projected on *ψuv*-space. **b/c**. Slow dynamics on 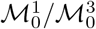, see system (3.53)/(3.56). Dark blue indicates the part of the orbit selected for the singular skeleton.

The idea behind this proof is similar to that of Theorem 3.3, in the sense that it is based on the existence of a special orbits 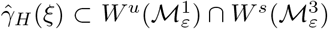 that take over the role of 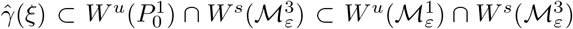 in the proof of Theorem 3.3 in combination with arguments based on the continuous dependence of initial conditions and on the symmetries in the system.

*Proof*. To show persistence, we first need to have a more precise characterization of the underlying skeleton. We choose a 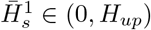 and define *ū*_*H*,min_ as the minimal value of the orbit 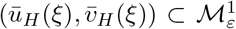 that follows the level set 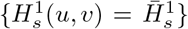 (3.54) (at leading order in *ε*), i.e. we set *ū*_*H*_ (0) = *ū*_*H*,min_ ≤ *ū*_*H*_ (*ξ*) for all *ξ* – where we note that (*ū*_*H*_ (*ξ*), 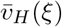) is periodic (see Fig. 3.8b). We define the exponentially short interval ℐ_*H*_ ⊂ {*z* = *v* = 0} by ℐ_*H*_ = {(*ψ, z, u, v*) = (*ψ*_0_, 0, *ū*_*H*,min_, 0)} with 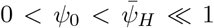 exponentially small and consider the two-dimensional manifold 𝒱_*H*_ spanned by the one-parameter family of solutions *γ*_*H*_ (*ξ*; *ψ*_0_) of (3.5) with initial conditions on ℐ_*H*_, i.e. *γ*_*H*_ (0; *ψ*_0_) = (*ψ*_0_, 0, *ū*_*H*,min_, 0). Note that the manifold 𝒱_*H*_ takes over the role played by 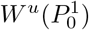 in the proof of Theorem 3.3.

Clearly, 𝒱_*H*_ is exponentially close to 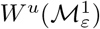] and since 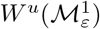 intersects 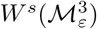 transversally it follows that 𝒱_*H*_ and 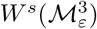 also intersect and that 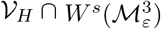 is one-dimensional. Since the orbits of (3.5) on 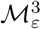 close to the level set 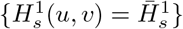 are periodic, we cannot assume that 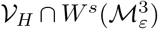 consists of only one orbit 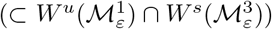: an orbit 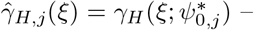 with 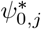 expo-nentially small – may first follow the periodic orbit on 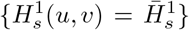 for *j* ≥ 0 circuits before it takes off from 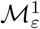 – necessarily near the jumping/take off point determined by 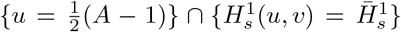. We define the orbit 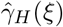 by 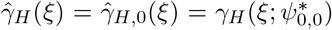, i.e. the orbit that does not make any additional circuits around the closed orbit level set 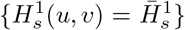, but ‘jumps away’ from 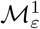 as soon as it crosses through 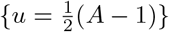.

By construction, the behavior of 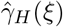 is similar to that of 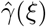 considered in the proof of Theorem 3.3: as it passes through 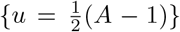, it takes off from 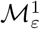 and, after its brief jump through the fast field, touches down on 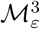 and subsequently follows the slow flow on 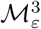 (at leading order determined by the level curve 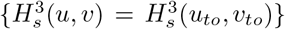 of (3.56)). Since 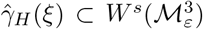 it does not take off again from 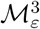. Equivalently, after touching down on 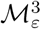, it does not intersect the plane {*z* = 0}. However, it does approach {*z* = 0} for *ξ* → ∞. Moreover, 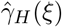 separates 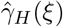 in two pieces (locally, near 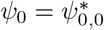): a part 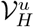 that does not cross through {*z* = 0} and a part 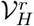 that does, i.e. the orbits 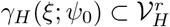 do intersect {*z* = 0} – say at *ξ* = *ξ*^*†*^(*ψ*_0_) – and return back to a neighborhood of 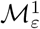 after that. Moreover, 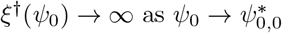 (as in the proof of Theorem 3.3).

Again as in the proof of Theorem 3.3: orbits 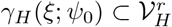 with *ψ*_0_ sufficiently close to 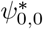 will follow 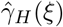along 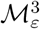 and thus cross through {*v* = 0} at a certain *ξ* = *ξ*^∗^(*ψ*_0_) *< ξ*^*†*^(*ψ*_0_). However, for *ψ*_0_ further away from 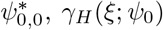 will take off from 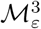 before it reaches {*v* = 0}: *ξ*^*†*^(*ψ*_0_) *< ξ*^∗^(*ψ*_0_). By continuity this (again) means that there must be a 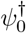 such that 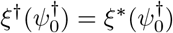.

The orbit 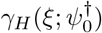 thus has both 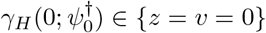 and 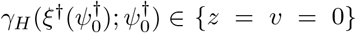. It follows by applying the reflectional symmetry (3.60) at both (end)points that 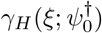 is a periodic orbit of (3.5) (with period 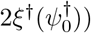 that by construction merges with the skeleton this proof started out with. □

#### 3.4.7 Geometric consistency of the 2-front periodic orbits on 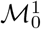 **and** 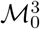

In Theorem 3.4, we proved the existence of 2-front periodic orbits that jump from 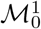 to 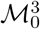 and back, in system (3.5). In this section, we investigate the observability of these periodic orbits, using geometric consistency condition (3.17). To this end, we want to use similar tools as in Section 3.3.2. Throughout this section, we use the same assumption as in Theorem 3.4, specifically that 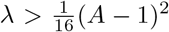. First we will perform the calculations for the system with the piecewise smooth function *f* (*K*), but in Lemma 3.5, we will extend the conclusions for the system with regularized function 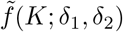.

To find the integral 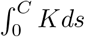 to leading order in *ε* and *δ*_1,2_, we compute it for the singular skeleton; only considering the parts on slow manifolds 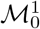 and 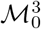. The dynamics on 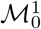 are described by system (3.53) and the dynamics on 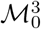 are described by system (3.56). For the slow dynamics of the singular skeleton, an orbit needs to connect in the jumping point 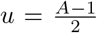, see Figure for a graphical representation of these orbits 3.9. To simplify computation, we derive one conserved quantity by evaluating the Hamiltonian on 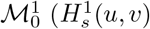 (3.54)) and Hamiltonian on 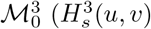, see (3.57)) in the jumping point 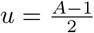 ;

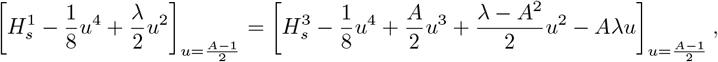

that can be reduced to

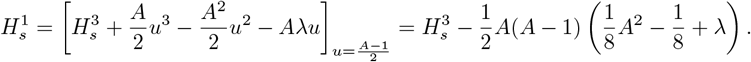

Now we can derive one conserved quantity 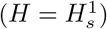,

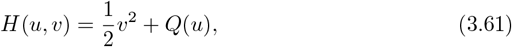

that connects slow orbits on slow manifolds 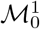and 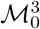 in the jumping point, where

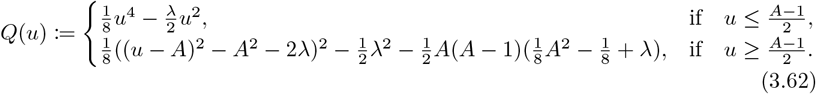

Note that there are two qualitatively different types of periodic orbits in the family of Theorem 3.4, as shown in Figure 3.9. The difference between these two orbits is in the location of steady state 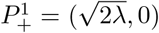 with respect to 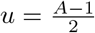. In Figure 3.9a, the orbit part on 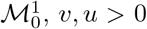 only increases. In Figure 3.9b however, the orbit on 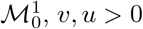 first increases but then decrease when *u* passes through 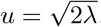. Note that for both these orbits Hamiltonian *H <* 0 (3.61)

**Figure 3.9:**
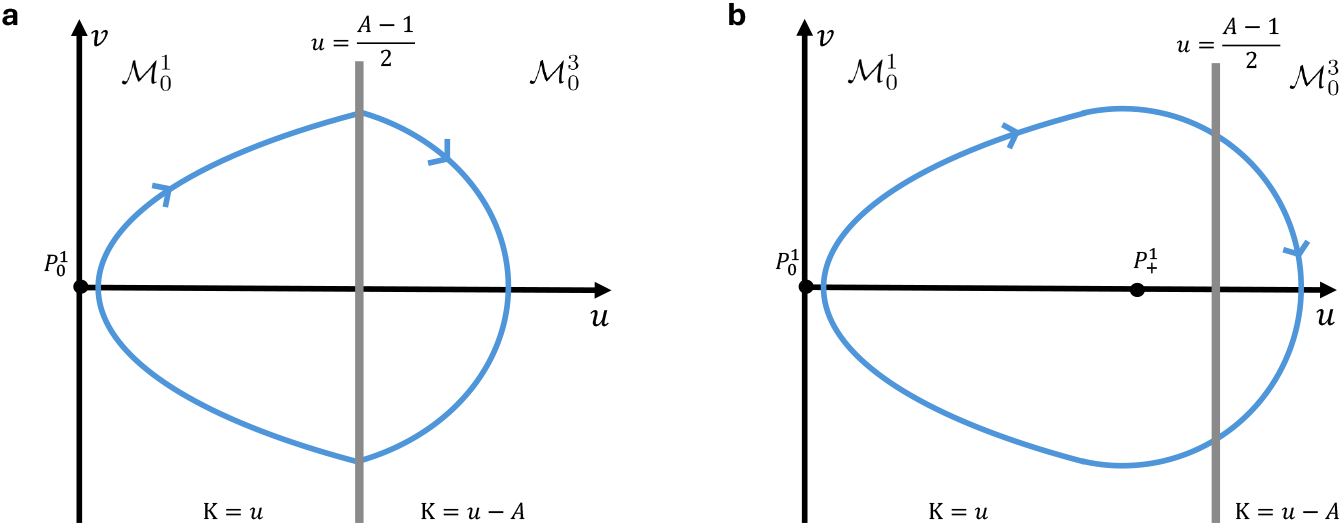
The slow flow of the 2-front periodic orbits from 3.4, projected on the *uv*-plane. **a**. 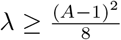 **b**. 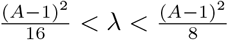

Similar to Section 3.3.2, we define *ū*_*p*_ (*ū*_*n*_) to be the unique value where the orbit intersects the *u*-axis for 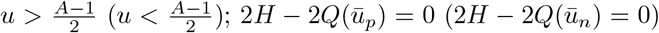. Moreover, let *Ĉ* be the period of the orbit. As before, we can find a relationship between *Ĉ* and the length of the curve *C*; there exists a number *n* ∈ ℕ such that *C* = *n · Ĉ*. We assume that 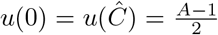 and *v*(0) = *v*(*Ĉ*) *>* 0. We define *B* by the unique value where 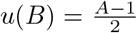 and 0 *< B < Ĉ*. By symmetry, 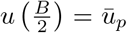 and 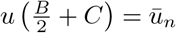.

We use the definition of *u* (3.2), *ψ* (3.2) and *K*_0_ (2.23) together with the location of slow manifold 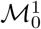 (3.23) and 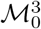 (3.25) to obtain the relationship between *u* and curvature *K*,

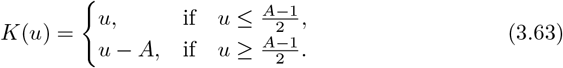

Now, we can split the orbits in two parts, 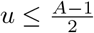 and 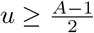,

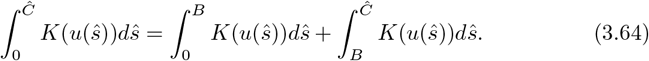

From the Hamiltonian (3.61) and the differential equations of *u* on 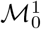(3.53) and on 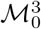(3.56), we derive

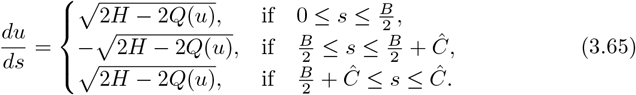

First we compute the second integral of (3.64) for 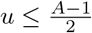:

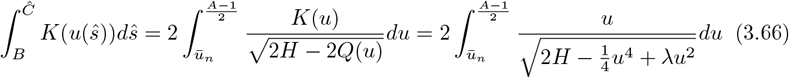

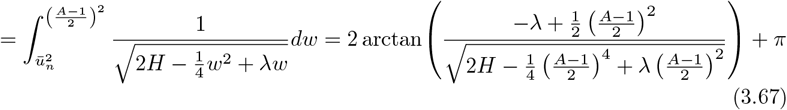

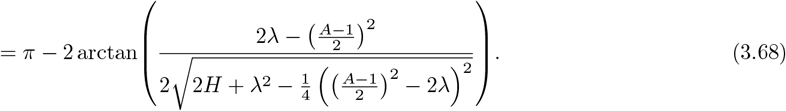

In the first equality of (3.66), we use the symmetry of the system and the derivative of *v*, (3.65). Furthermore, we substitute *K*(*u*) (3.63) and *Q*(*u*) (3.62). To obtain (3.67), we take *w* := *u*^2^. In Section 3.3.2, we derived the integral of (3.67), see equation (3.40). Note that,

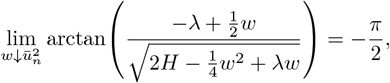

since 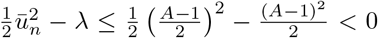. Lastly, in (3.68), we rewrite the argument of the arctan and we use that arctan(−*x*) = − arctan(*x*).

Now, we compute the first integral of equation (3.64) (for 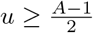):

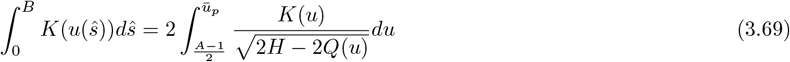

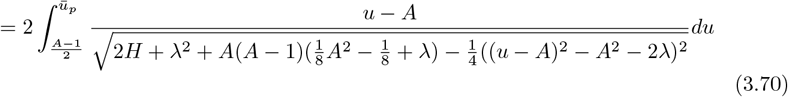

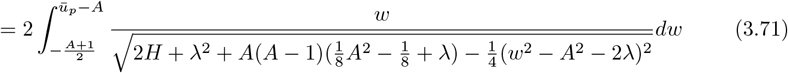

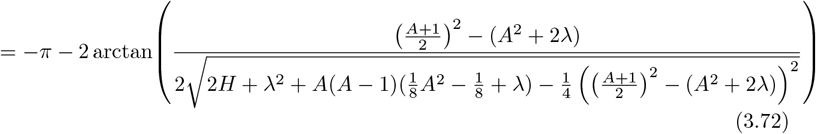

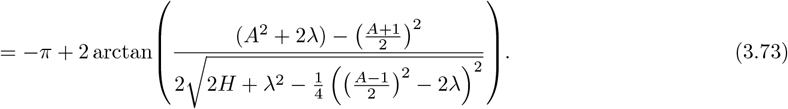

In (3.70) we use symmetry, 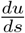 (3.65) and the definition of function *Q*(*u*) (3.62). In (3.71) we substitute *w* := *u* − *A*. In (3.72), we use that

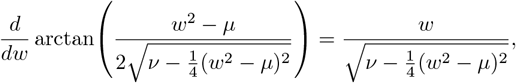

where 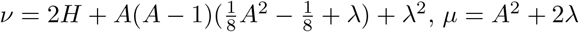 and

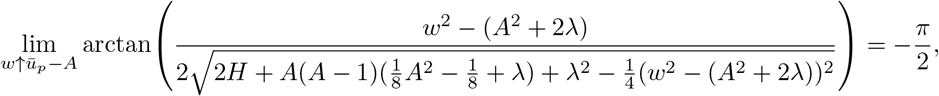

since 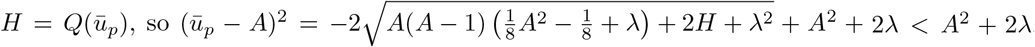. Lastly, in (3.73), we use that arctan(−*x*) = −arctan(*x*) and that 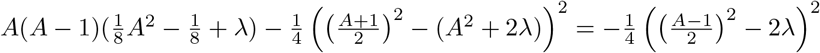

We use the above computations to prove Lemma 3.5.

##### Lemma 3.5.

*Let* 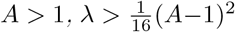 *and let* 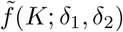 *be a smooth regularization of f* (*K*) (2.26). *Consider periodic orbit* 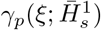*(see Theorem 3*.*4). For δ*_1,2_ *and ε sufficiently small*, 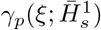 *does not meet geometric consistency condition* (3.17) *and is therefore, not observable as a solution of the PDE* (1.4).

*Proof*. In this proof we use similar arguments as in the proof of Lemma 3.2. By rewriting (3.68) and (3.73), we can deduce

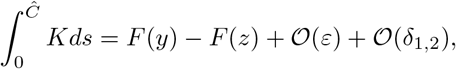

for *F* (*x*) := 2 arctan(*x*) and 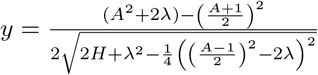 and 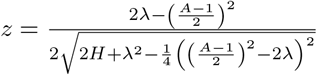

Since *F* (*x*) is strictly increasing, *F* (*y*) = *F* (*z*) if and only if 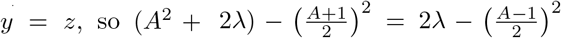, which is only true when *A* = 1. However, since *A >* 1 for the orbits we are investigating, *y* ≠ *z* for these periodic orbits. However, if we assume that *y*≠ *z*, we can take *ε* and *δ*_1,2_ small enough such that *F* (*y*) − *F* (*z*) + *𝒪* (*ε*) + *𝒪* (*δ*_1,2_) ≠ 0.

See Figure 3.10 for a plot of 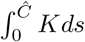 in leading order of *ε* and *δ* over *λ*_1,2_ for some values of *A* and *H*. Note that the integral increases for a small *λ* interval, whenceforth it decreases rapidly. Observe that for fixed *λ*, the integral decreases when we decrease *A*. We can interpret the results of Lemma 3.5 as follows. For an orbit (which describes the curvature of a curve over the curvilinear coordinate) to be observable, it needs to have “as much negative curvature as positive curvature”. The part of the orbit on 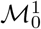 describes the curve when no morphogen is present. Here, the curve has positive curvature (is curving outward). The part of the orbit on 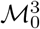describes the curve when there is morphogen present. Here, the curvature is negative. The analysis shows that the dynamics on and the geometry of the slow manifold make it impossible for the total amount of negative curvature to be enough, i.e. to start and end with the same slope of the curve (periodic boundary conditions), if we fix *λ* and the order one approximation of the periodic orbit. Note, however, that if we fix *δ*_1,2_ and *ε*, there might still be an observable periodic orbit. We furhter elaborate on this topic in Section 3.5

**Figure 3.10:**
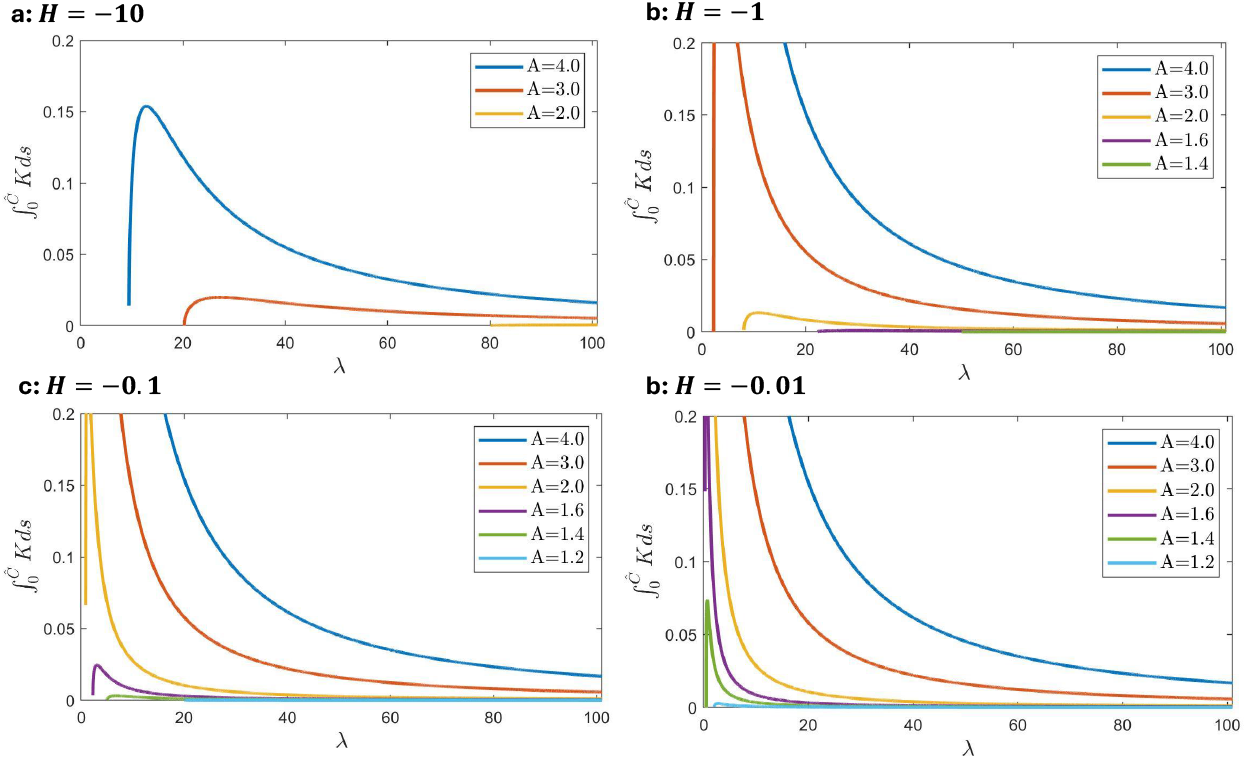
The integral of the curvature in leading order of *ε* and *δ*_1,2_ plotted over values of the Lagrange multiplier *λ*.

### 3.5 The possibility of geometric consistent periodic orbits

In Sections 3.3 and 3.4 we study the possible periodic orbits in system (3.6) and their geometric consistency with respect to the evolution equation (1.4). In the weak and strong interplay regime, we prove that periodic orbits (uniquely) exist for *ε* and *δ*_1,2_ small enough, see Theorems 3.1 and 3.4. However, these orbits do not comply with the geometric consistency conditions when *ε* and *δ*_1,2_ are sufficiently small, see Lemmas 3.2 and 3.5. Still, it might be possible for a periodic orbit to be observable in PDE (3.6), for the following reason. In Lemmas 3.2 and 3.5 we first fix *H* and *λ* and then take *δ*_1,2_ and *ε* small. However for the flow of the PDE, this process is reversed: first the parameters *δ*_1,2_ and *ε* are fixed, and then the PDE converges to a stationary spatially periodic solution, selecting *H* and *λ*. This allows for the possibility to take *δ*_1,2_ and *ε* small enough for a stationary, spatially periodic solution to exist (Theorems 3.1/3.4), but not so small that the solution becomes unobservable.

In Figure 3.3, for the weak interplay regime, we observe that the (leading order expression for the) value of 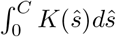 decreases when *H* decreases or −*λ* increases. This is an indication that when *δ*_1,2_ and *ε* are small, a larger −*λ* is selected. Figure 3.10 shows a similar response for the strong interplay regime: when *λ* increases, the leading order value of 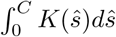 decreases. Now note that the analysis leading to Lemmas 3.2 and 3.5 is based on the 𝒪 (1) properties of the constructed patterns, i.e. their singular skeletons. Thus a more accurate derivation of integral (3.17) over the curvature will have an 𝒪 (*δ*_1,2_, *ε*) correction. As the plots in Figures 3.3 and 3.10 suggest, consistency condition may be made to hold for corrected, more accurate approximation of (3.17) by choosing *λ* sufficiently large.

Furthermore, in this parameter regime, a larger value of *A* corresponds to a larger value of the integral over the curvature, suggesting that for larger values of *A*, it might be more difficult for a pattern to be observable. We further investigate these aspects in the next section, where we perform a numerical simulation.

## 4 Numerical simulations

In Section 3, the steady state phase space of system (1.4) is analyzed. The analysis provides possible steady state solutions; however, it does not indicate which solutions the model converges to. For observability, the solutions must be stable, and the initial condition must be in the basin of attraction of this stable steady state. Furthermore, the existence analysis focused on solutions in a specific parameter regime, namely *ε* small enough and all other parameters of 𝒪 (1).

To further analyze the possible steady state solutions of system (1.4) and their temporal stability, we use numerical simulation. For implementation, space discretization and finite difference are used to approximate spatial derivatives. Moreover, the TR-BDF2 method is used to simulate the evolution of system (1.4) over time [1]. This method regulates the value of the Lagrange multiplier *λ* to keep the length of the curve, *C*, constant over time, see equation (2.10). The complete code, based on the work of Kazarnikov et. al. [21], can be found at https://github.com/DaphnN/Mechanochemicalmodel.

As we show in this section, the results of our numerical simulations show a rich spectrum of possible stable steady state solutions. Below, we display some of these possible solutions and demonstrate how they fit into the analytical framework. We follow the case separation from Section 3, distinguishing weak and strong morphogen-curvature interplay, quantified by the parameter *A*, see equation (3.7). For the curvature dependent morphogen production, we use the smooth regularization 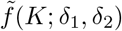, as defined in (2.27).

### 4.1 Simulations in the weak morphogen-curvature interplay regime, *A <* 1

Here the simulation results in the weak morphogen-curvature interplay regime are presented. Figure 4.1a shows the stable steady state solution when simulating the PDEs, system (1.4), for *A* = 0.5, *ε* = 0.01, *L* = 3, *C* ≈ 3.0021, *δ*_1,2_ = 0.01. In this parameter regime, the solution for the height of the curve, *h*, is slowly changing in space. The morphogen concentration, *ϕ*, attains a single pulse solution; the concentration is zero outside an 𝒪 (1) region. In Figure 4.1b, the corresponding curvature, *K* (2.6), of the steady state curve is shown. Note that in the positive concentration region, the curvature of the curve is negative. Also note that the derivative of the curvature *K* changes abruptly at the boundaries of the morphogen pulse.

The simulation is terminated when the shape of the solution remains constant over time. However, aslow horizontal shift of the solution could be observed. Since periodic boundary conditions (see Section 2.4) allow for a horizontal translation, any solution to the PDE system (1.4) comes in a family of horizontal translates. Hence, it is likely that the numerically observed horizontal drift is caused by numerical solutions that are close to a stable steady state but not equal to it. For the exact steady state solution, the eigenvalue in the direction of the equivalent, shifted solutions equals zero. However, for a steady state solutions with a numerical error this eigenvalues is small but not equal to zero, causing a slow horizontal drift.

The numerical solution shown in Figure 4.1a can be compared with the analytical insights obtained in Section 3.3. Figure 3.1 shows the shape of the invariant manifolds for *A <* 1 in *ψu*-space. To visualize the correspondence of the simulation to the analysis, the numerical steady state solution, presented in Figure 4.1a, is plotted in *ψuv*-space, together with the invariant manifold for *δ*_1,2_ = 0, see Figure 4.1c. As expected from the analysis, the solution curve is close to the *δ*_1,2_ = 0 invariant manifold. The existence of periodic orbits of the type shown in Figure 4.1c was proven in Theorem 3.1. Furthermore we proved that a particular periodic orbit (for fixed *H* and *λ*) did not meet geometric consistency condition (3.1), 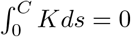, for *δ*_*1,2*_ and *ε* small enough, Lemma 3.2. However, Section 3.5 explained that by first fixing *δ*_1,2_ and *ε* small, an orbit and Lagrange multiplier *λ* might be selected that can compensate with its higher order terms for the first order approximation of the integral over the curvature, see Figure 3.3. Since 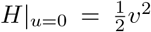 we can use the intersection of the periodic orbit in Figure 4.1c with the *v*-axis to find *H* ≈ 0.014. We find that 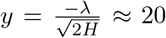, which corresponds to a very small first-order approximation of the integral over the curvature, see Figure 3.3. Therefore, the small parameters *ε* and *δ*_1,2_ are in this case not ‘sufficiently small’ to conclude non-observability of the analytical solution. In other words, a large enough value for parameters*ε, δ*_1,2_ *>* 0 possibly combined with the finite number of points (*N* = 256) representing a continuous curve introduce sufficient freedom and possibly numerical inaccuracy to allow this numerical solution to be a steady state solution (3.3) but also obey the geometric consistency condition (3.17).

**Figure 4.1:**
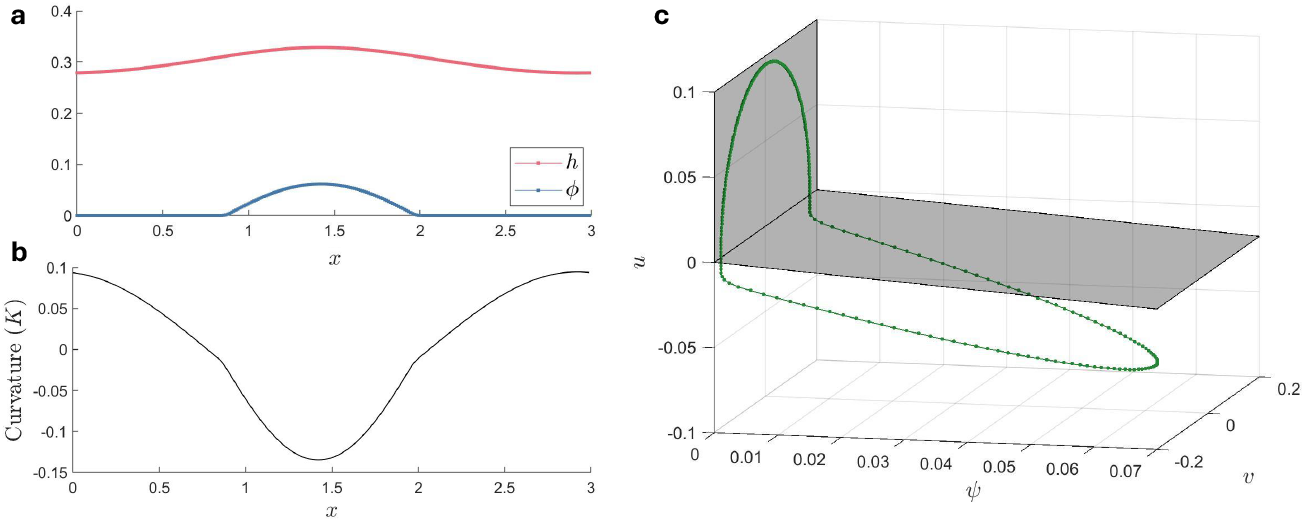
The numerically obtained steady state solution of system (1.4) for *A* = 0.5 (*β* = 1, *α* = 2 and *η* = 1), *ε* = 0.01 (*D* = 10^−4^), *L* = 3, *C* = 3.0021, *δ*_1,2_ = 0.01 and number of discretization points, *N* = 256. The Lagrange multiplier for this solution, *λ* ≈ − 3.34. **a**. Steady state solution of the height of the curve, *h*(*x*), and concentration of morphogen, *ϕ*(*x*). **b**. Plot of *K* (2.6), the curvature of the steady state solution curve, *h*(*x*), of Figure a. **c**. Steady state solution plotted as an orbit in *ψuv*-space.

To investigate the influence of the curve length on the type of steady state solutions found, we run simulations with a range of curve lengths *C*. Figure 4.2a shows the steady state solutions plotted in *ψuv*-space. These orbits correspond to different values of the Lagrange multiplier *λ* see Figure 4.2c. Therefore, the steady state solutions live in different phase spaces; see equation (3.3) for the influence of the value of *λ* on the dynamics. In Figure 4.2c, the steady state solution curve *h*(*x*) and morphogen concentration *ϕ*(*x*) are plotted for the largest and smallest curve length value *C*. Observe that there is a larger curve deformation when the curve is longer (*C* = 3.026). This is expected from the geometric limitations of the curve; the curve length cannot change with respect to the domain length *L*. These larger curve deformations correspond to a higher pulse solution for *ϕ*.

In Figure 4.3a, the numerical steady state solutions are presented for different values of *A <* 1. Note that the slow manifold shape and location change for the *ψ >* 0 region, as expected from the analysis. This results in a more steep pulse solution when *A* is larger; see Figure 4.3b. Interestingly, the relationship between the value of *A* and the Lagrange multiplier *λ* corresponding to the steady state solution for that value of *A* seems to be linear. As expected from Figure 3.3, when we increase 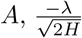 decreases.

**Figure 4.2:**
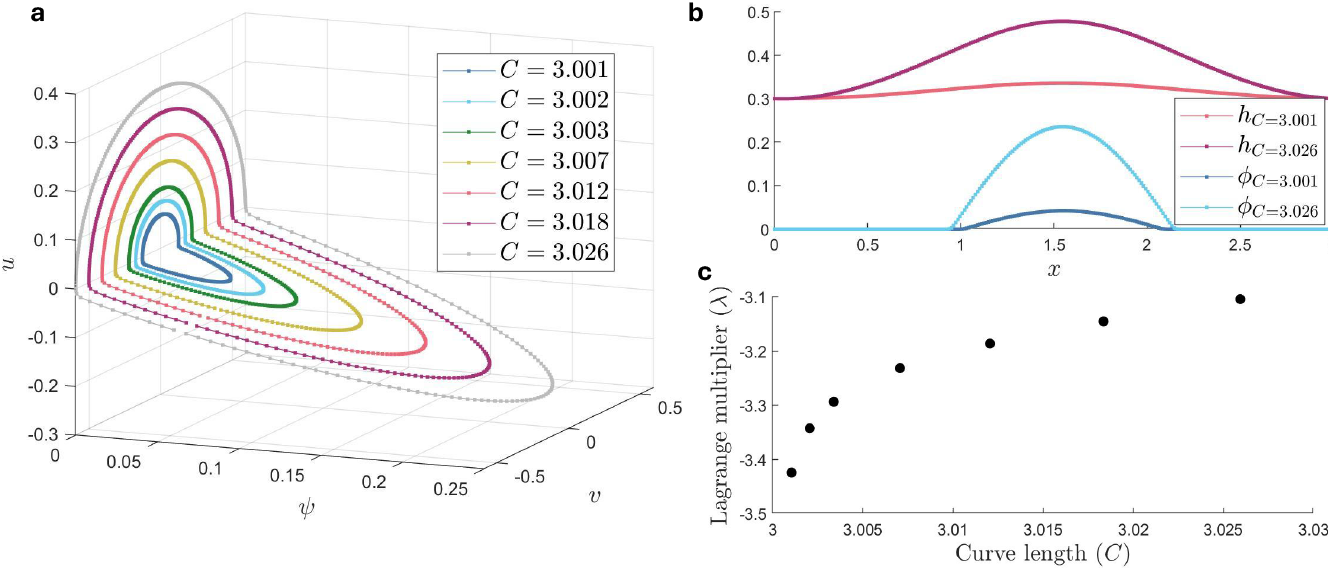
The numerically obtained steady state solution of system (1.4) for a range of curve length values, *C*. For the other parameters *A* = 0.5 (*β* = 1, *α* = 2 and *η* = 1), *ε* = 0.01 (*D* = 10^−4^), *L* = 3, *δ*_1,2_ = 0.01 and number of discretization points, *N* = 256. **a**. Steady state solutions plotted as orbits in *ψuv*-space. **b**. Steady state solutions of the height of the curve, *h*(*x*), and concentration of morphogen, *ϕ*(*x*) for *C* = 3.001 and *C* = 3.026. **c**. Lagrange multiplier values, *λ*, corresponding to the steady state solutions plotted in Figure a.

To study the influence of small parameters, we run simulations for a range of *ε* and *δ*_1,2_. Figure 4.4 shows the numerical steady state solution, plotted in *ψuv*-space, when decreasing *ε*. Note that these solutions are approaching the slow manifolds derived in Section 3.2, when *ε* → 0. Moreover, the value of the Lagrange multiplier, *λ*, remains approximately constant. Figure 4.5 shows the steady state solutions for decreasing values of *δ*_1,2_. In Figure 4.5a, we observe that the orbit is getting closer and closer to the slow manifold for *δ*_1,2_ = 0, as plotted in Figure 4.1c.

Overall, we see that for the parameter settings used in this section, the value of the Lagrange multiplier was always negative. Furthermore, we do not observe any steady solution orbit that lies on the *ψ* = *A* part of the slow-invariant manifold.

### 4.2 Simulations in the strong morphogen-curvature interplay regime, *A >* 1

In this section we show the simulation results of system (1.4) for the strong interplay regime, *A >* 1. From the analysis of the slow manifolds of the steady state phase space, Section 3.4, we expect to find very different steady state solutions here compared to the weak interplay regime. In Theorem 3.4 we proved the existence of two-front periodic solutions, jumping from slow manifold 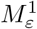 to 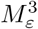. However, we proved that a periodic orbit did not comply with geometric consistency condition (3.1),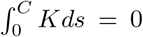, for *δ* _1,2_ and *ε* small enough, see Lemma 3.5. In Section 3.5, however, we explained that by first fixing *δ*_1,2_ and *ε* small, an orbit and Lagrange multiplier *λ* might be selected that can compensate for the first order approximation of the integral over the curvature with its higher order terms in *δ*_1,2_ and *ε*.

In Figure 4.6, we plot the numerically obtained steady state solution for *A* = 2, *ε* = 0.01, *L* = 3, *C* = 3.0021, *δ*_1,2_ = 0.01. Figure 4.6a shows the height of the curve *h*(*x*) and the morphogen concentration *ϕ*(*x*) in steady state. We observe a slow vertical shift of the curve *h*(*x*), when the solution is in steady state. This is likely caused by the same phenomenon as the horizontal shift for *A <* 1. Namely, all vertical translates are part of the same family of solutions of PDE (1.4), hence the corresponding zero eigenvalue can cause a slow vertical drift in numerically obtained stable steady state solutions.

**Figure 4.3:**
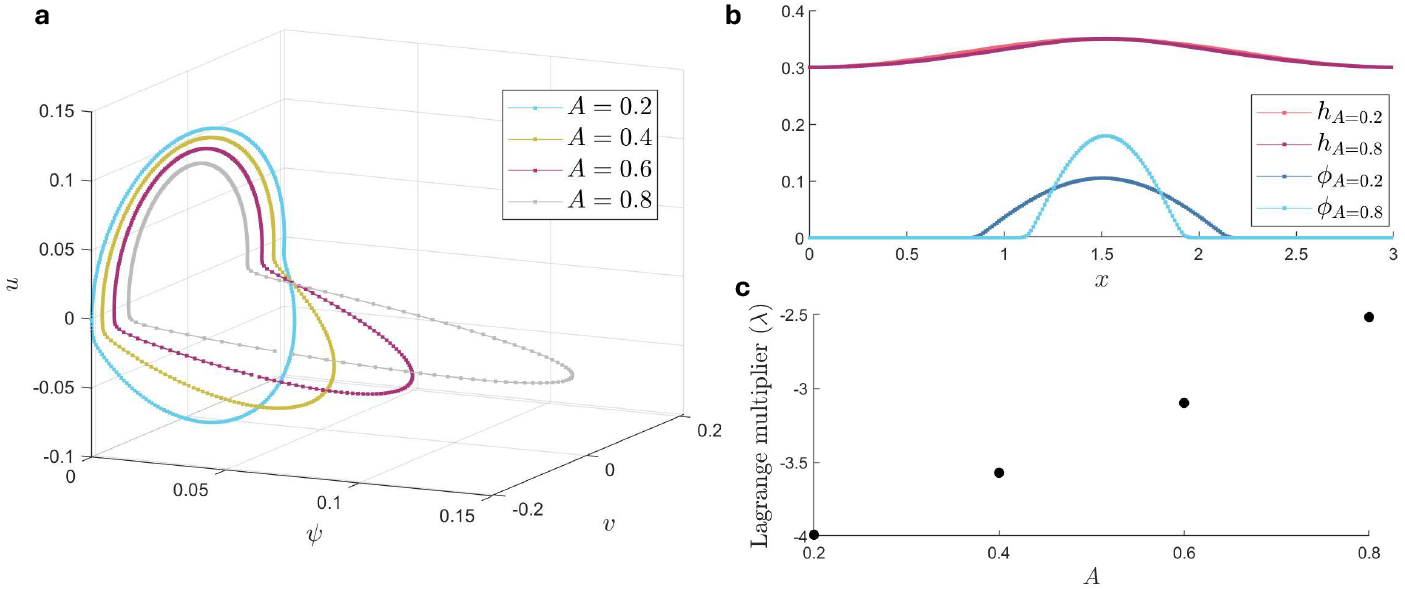
The numerically obtained steady state solution of system (1.4) for a range of *A* values, by changing *β* but keeping *η* = 1 and *α* = 1 constant. For the other parameters *ε* = 0.01 (*D* = 10^−4^), *L* = 3, *C* = 3.0021, *δ*_1,2_ = 0.01 and number of discretization points, *N* = 256. **a**. Steady state solutions plotted as orbits in *ψuv*-space. **b**. Steady state solutions of the height of the curve, *h*(*x*), and concentration of morphogen, *ϕ*(*x*) for *A* = 0.2 and *A* = 0.8. **c**. Lagrange multiplier values, *λ*, corresponding to the steady state solutions plotted in Figure a.

**Figure 4.4:**
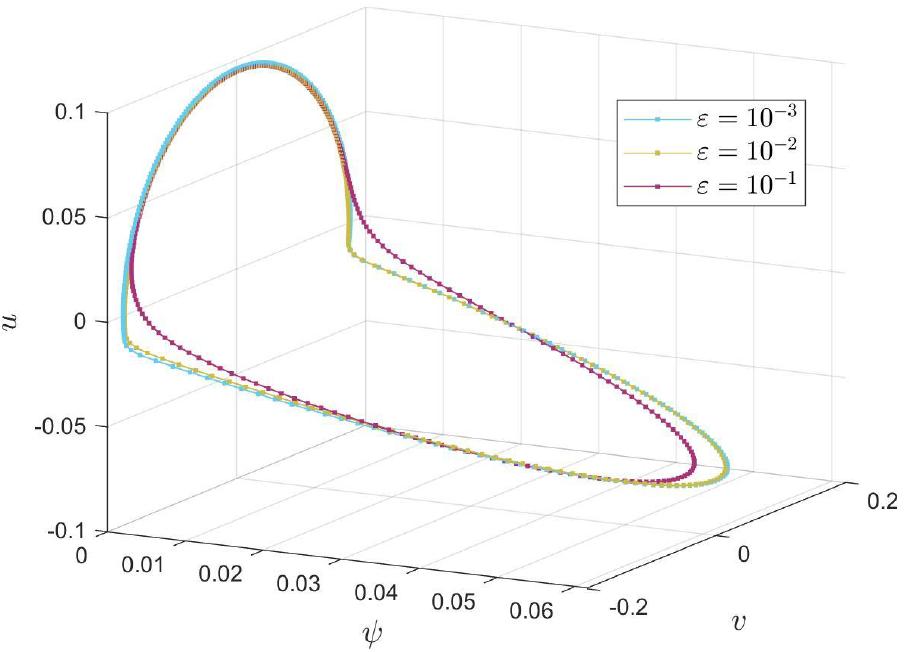
The numerically obtained steady state solution of system (1.4) for changing value of *ε*. For the other parameters; *A* = 0.5 (*β* = 1, *α* = 2 and *η* = 1), *L* = 3, *C* ≈ 3.0021, *δ*_1,2_ = 0.01 and number of discretization points, *N* = 256. The Lagrange multiplier is approximately the same for these three simulations: 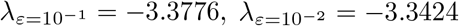 and 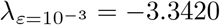

**Figure 4.5:**
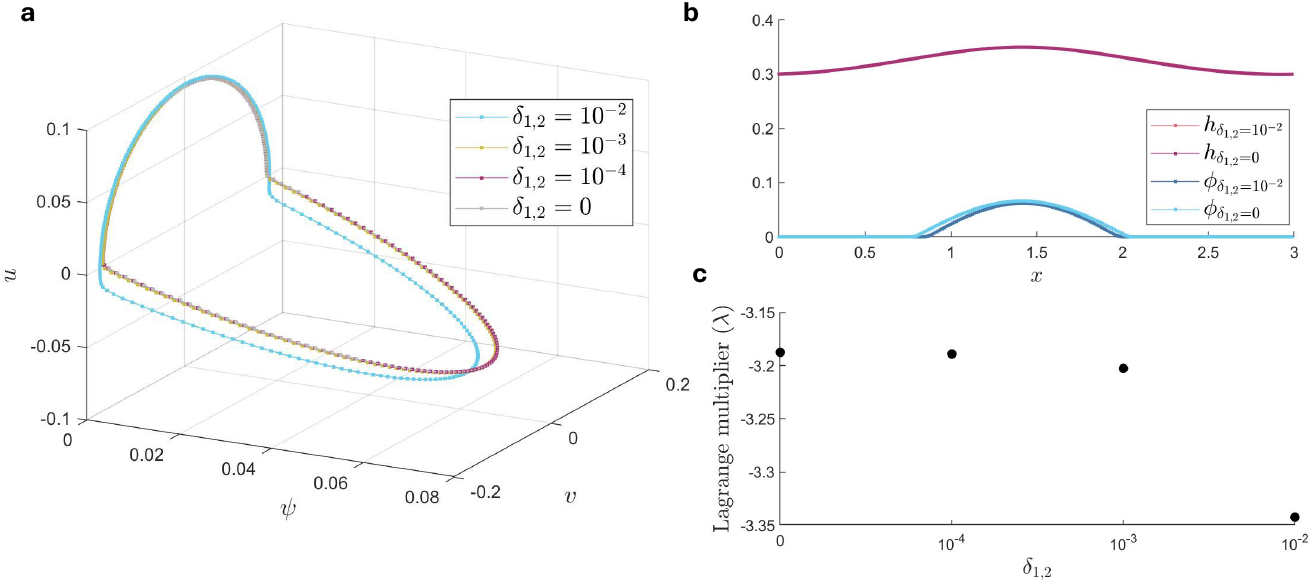
The numerically obtained steady state solution of system (1.4) for a range *δ*_1,2_ values. For the other parameters *A* = 0.5 (*β* = 1, *α* = 2 and *η* = 1), *ε* = 0.01 (*D* = 10^−4^), *L* = 3, *C* = 3.0021, and number of discretization points, *N* = 256. **a**. Steady state solutions plotted as orbits in *ψuv*-space. **b**. Steady state solutions of the height of the curve, *h*(*x*), and concentration of morphogen, *ϕ*(*x*) for *δ*_1,2_ = 0.01 and *δ*_1,2_ = 0. **c**. Lagrange multiplier values, *λ*, corresponding to the steady state solutions plotted in Figure a.

Figure 4.6a displays a 2-front solution for the morphogen *ϕ*(*x*), however, the curve *h* does not have rapid jumps. If we consider the curvature of *h*(*x*), exhibited in Figure 4.6b, we do see that there are jumps in the same location as *ϕ*. Note that *h* is curving outward when morphogen is present, as expected.

To further compare the numerical solutions with the analysis in Section 3, we plot the steady state solution in *ψuv*-space together with the normally hyperbolic slow manifolds for 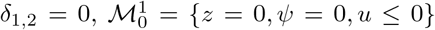 and 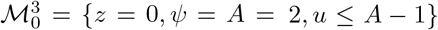. Note that this orbit is approximately following the singular skeleton of the 2-front periodic orbit from Theorem 3.4. The orbit consists of a slow part on the trivial slow manifold, 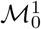and a slow part on 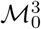. Observe that the quick jump occurs approximately when 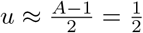, as expected.

Figure 4.7 shows the numerical steady state solutions for different curve lengths *C*, (2.10). When we plot these orbits in *ψuv*-space (Figure 4.7a), we see that the orbit is less symmetric for higher values of *C*. These asymmetric orbits do not correspond to Theorem 3.4, where we actually use the symmetry of the system to prove the existence of two front orbits. Interestingly, the Lagrange multiplier *λ* is quite large for (more) symetric orbits (with curve length *C* closer to *L* = 3), see Figure 4.7c. This might be caused by the first order approximation of the integral over de curvature, see Figure 3.10. Observe that the integral over the curvature increases for a very small interval of Lagrange multiplier *λ*, negligible for numerical simulations. Furthermore, Figure 3.10 shows that a larger Lagrange multiplier *λ* corresponds to a smaller first order approximation of the integral of the curvature. This is possibly the reason for a larger Lagrange multiplier; the higher order terms (in *δ*_1,2_ and *ε*) might be able to compensate for the non-zero first order approximation of the integral over the curvature. We should however question the validity of the analysis in this parameter regime. If the value of the Lagrange multiplier *λ* increases as e.g. 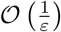, the analysis is no longer valid. In Figure 4.7b, we plot the curve *h*(*x*) and the morphogen concentration *ϕ*(*x*) corresponding to the orbits in Figure 4.7a with the longest and shortest curve. Note that when *C* is higher, the deformations are larger and the pulse solution is wider. This is an expected phenomenon from the geometry (assuming that there is only one pulse), since *C* cannot change compared to *L* is this model.

**Figure 4.6:**
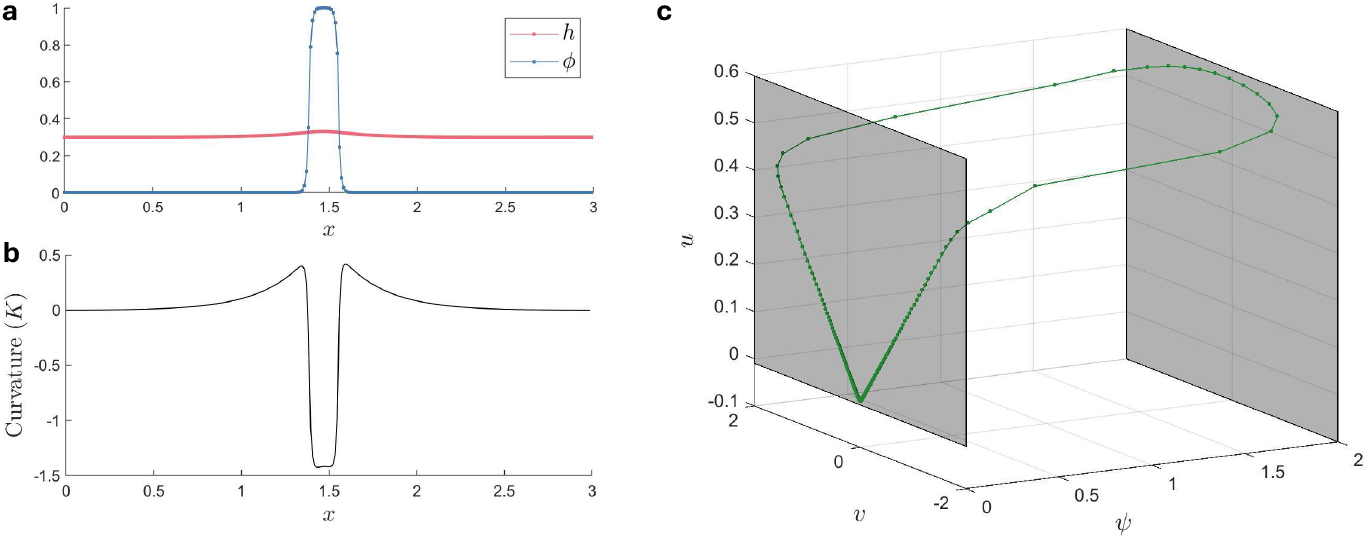
The numerically obtained steady state solution of system (1.4) for *A* = 2 (*β* = 2, *α* = 1 and *η* = 1), *ε* = 0.01 (*D* = 10^−4^), *L* = 3, *C* = 3.0021, *δ*_1,2_ = 0.01 and number of discretization points, *N* = 256. The Lagrange multiplier for this solution, *λ ≈* 15.48. **a**. Steady state solution of the height of the curve, *h*(*x*), and concentration of morphogen, *ϕ*(*x*). **b**. Plot of *K* (2.6), the curvature of the steady state solution curve, *h*(*x*), of Figure a. **c**. Steady state solution plotted as an orbit in *ψuv*-space. The pink planes correspond to the normally hyperbolic slow manifolds, 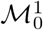 and 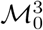.

**Figure 4.7:**
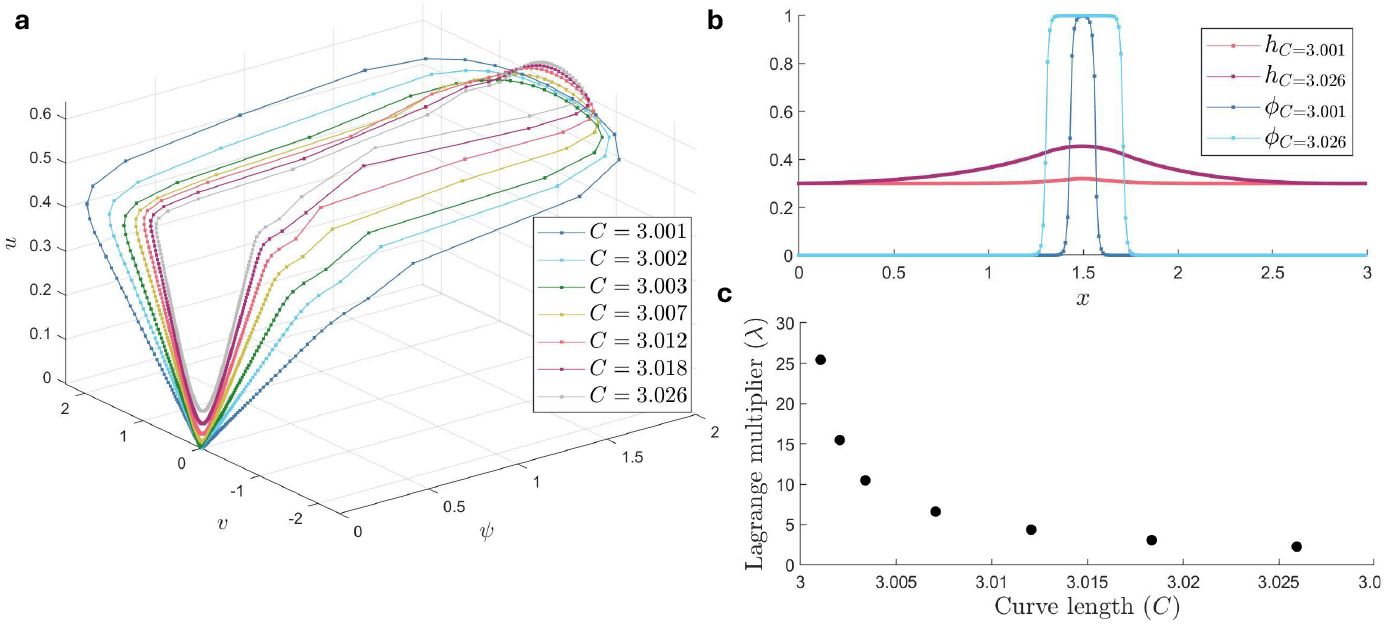
The numerically obtained steady state solution of system (1.4) for a range of curve length values, *C*. For the other parameters *A* = 2 (*β* = 2, *α* = 1 and *η* = 1), *ε* = 0.01 (*D* = 10^−4^), *L* = 3, *δ*_1,2_ = 0.01 and number of discretization points, *N* = 256. **a**. Steady state solutions plotted as orbits in *ψuv*-space. **b**. Steady state solutions of the height of the curve, *h*(*x*), and concentration of morphogen, *ϕ*(*x*) for *C* = 3.001 and *C* = 3.026. **c**. Lagrange multiplier values, *λ*, corresponding to the steady state solutions plotted in Figure a.

**Figure 4.8:**
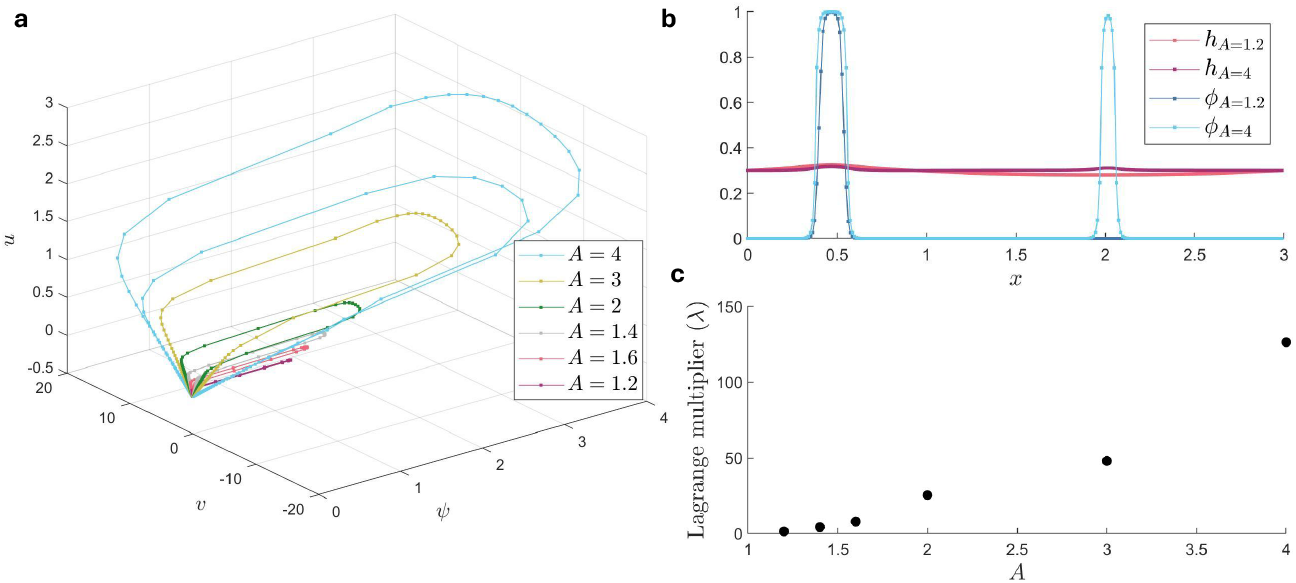
The numerically obtained steady state solution of system (1.4) for a range of *A* values, by changing *β* but keeping *η* = 1 and *α* = 1 constant. For the other parameters *ε* = 0.01 (*D* = 10^−4^), *L* = 3, *C* = 3.0021, *δ*_1,2_ = 0.01 and number of discretization points, *N* = 256. **a**. Steady state solutions plotted as orbits in *ψuv*-space. **b**. Steady state solutions of the height of the curve, *h*(*x*), and concentration of morphogen, *ϕ*(*x*) for *A* = 1.2 and *A* = 4. **c**. Lagrange multiplier values, *λ*, corresponding to the steady state solutions plotted in Figure (a).

We are also interested in the influence of parameter *A* on the steady state solutions, see Figure 4.8. As we expect from the location of the normally hyperbolic manifold 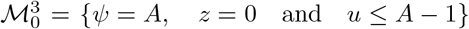 (3.25), the maximal *ψ* value of the periodic orbits increases when we increase *A*, see Figure 4.8a. To change *A*, we keep *α* and *γ* constant and only change *β*. Since we rescale 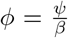, the hight of the front in *ϕ*, see Figure 4.8b, does stay constant. Figure 4.8c shows that Langrange multiplier *λ*, increases rapidly when we incrase *A*, which is expected from Figure 3.10; the larger *A*, the larger the first order approximation of the integral over the curvature. As mentioned above, *λ* scaling as, e.g. 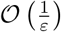 would greatly influence the analysis. This might be causing the solution with two different pulses that we observe when *A* = 4, related to a very large value of the Lagrange multiplier *λ*.

Lastly, we investigate the influence of small parameters *ε* and *δ*_1,2_ on the steady state solutions. Firstly, Figure 4.9 shows that when we take *ε* not small enough (*ε* = 0.1), the periodic orbit is not a 2-front solution. However, if we take *ε* too large (*ε* = 0.001), it makes abrupt jumps that do not correlate to a jump in the fast dynamics of the system. This might be caused by the geometric consistency condition as explained in Section 3.5. However, it could also be influenced by the low number of discretization points: we do not see any points halfway in the fast jump in *ψ*. In Figure 4.10, we display the influence of *δ*_1,2_ on the steady state solutions. Interestingly, when *δ*_1,2_ *≤*0.001, there are two two-front pulses. We also observe that the Lagrange multiplier *λ* increases. This behavior might be caused by another selection process as the periodic orbit needs to have the correct arc length *C* and domain length *L*, but could also be caused by the finite number of discretization points used in the simulation.

**Figure 4.9:**
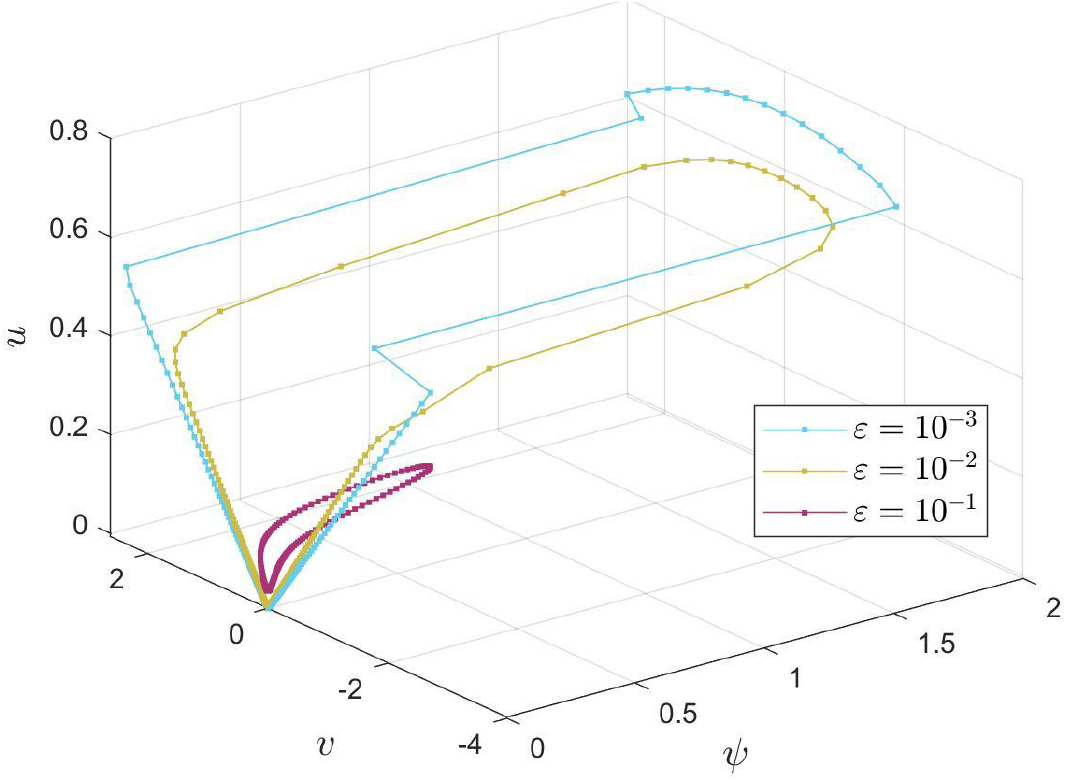
The numerically obtained steady state solution of system (1.4) for changing value of *ε*. For the other parameters; *A* = 2 (*β* = 2, *α* = 1 and *η* = 1), *L* = 3, *C* ≈ 3.0021, *δ*_1,2_ = 0.01 and number of discretization points, *N* = 256.

**Figure 4.10:**
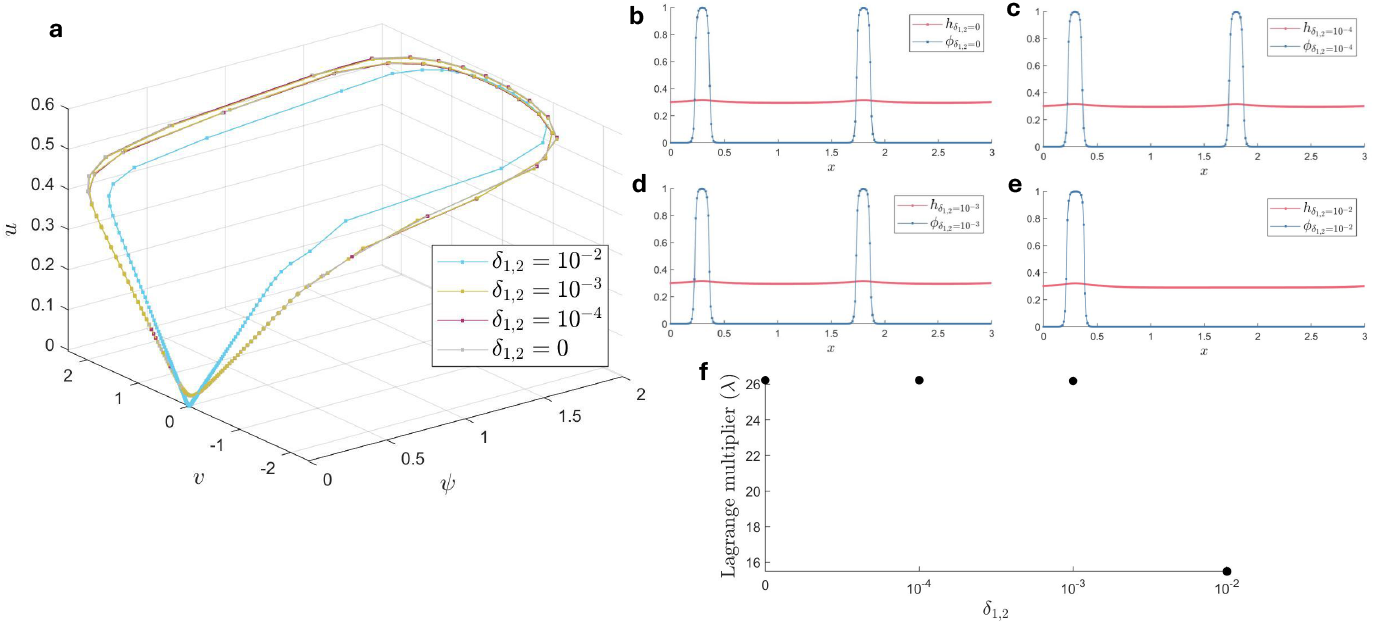
The numerically obtained steady state solution of system (1.4) for a range *δ*_1,2_ values. For the other parameters *A* = 2 (*β* = 2, *α* = 1 and *η* = 1), *ε* = 0.01 (*D* = 10^−4^), *L* = 3, *C* = 3.0021, and number of discretization points, *N* = 256. **a**. Steady state solutions plotted as orbits in *ψuv*-space. **b-e**. Steady state solution of the height of the curve, *h*(*x*), and concentration of morphogen, *ϕ*(*x*) corresponding to the curves in figure a. **f**. Lagrange multiplier values, *λ*, corresponding to the steady state solutions plotted in Figure (a).

## 5 Discussion

In this paper, we derive a mathematical model (1.4) that serves as a phenomenological mechanism for the formation of spatial inhomogeneities and symmetry breaking in morphogenesis. This model, based on the work of Mercker et al., describes the spread of a morphogen, interacting with the curvature of a biological surface, e.g. a cell membrane or the surface of a tissue [29]. To study the patterns that this model produces, we used a combination of analytical and numerical tools. In this section, we discuss the main results of this study and suggest some possibilities for follow-up research.

The mechanochemical model studied in this paper is a simplified description of realistic tissue surface or cell membrane dynamics. We assumed that the surface can be represented by a curve that is a graph of a function. Furthermore, the evolution equation of the curve (1.4) is derived by taking the *L*^2^-gradient flow of the Helfrich energy, introducing a time evolution that minimizes this energy over time. The Helfrich energy incorporates the difference between the local curvature, *K*, and the preferred local curvature, *K*_0_(Φ), given a certain morphogen concentration. For analytical convenience, we drop the variation of the curve in the tangential direction by setting the second part of equation (2.21) to zero. This simplified model can be used as a starting point for understanding the full model. The second term of equation (2.21) might have an influence on the type of solutions or patterns that the model shows, especially in relation to the geometric consistency conditions.

The dynamics of the morphogen are described by a reaction diffusion equation (1.4b) with a curvature-dependent morphogen production, *f* (*K*), see (2.26) and constant degradation, *α*. We assume that the morphogen is diffusing slowly, which reflects the observation that mechanistic changes in the tissue occur on a faster time scale than the diffusion of the morphogen over the tissue surface. Note that the choice made for the curvature-dependent production rate directly determines the shape of the critical manifold ℳ_0_ (3.21), and therefore determines the spatial structure of pattern solutions to (1.4).

Based on the geometric context of the mechanochemical model, we derived periodic boundary conditions in Section 2. These periodic boundary conditions allow the model to represent a closed curve to leading-order, when *h* represents deviations from a circle with a large radius. However, the small correction terms that are introduced when the graph of *h* is interpreted as the deviations of a large circle might be important in the analysis of the possible steady state solutions, especially in relation to the geometric consistency conditions, derived in Section 3.1. In other words, the “background curvature” that is neglected in our approach might influence which curves are observable.

The objective of Section 3 was to analyze which patterns could be observed in the model (1.4), using GSPT. Analyzing the invariant manifolds of the system of ODEs (3.3) showed that the type of possible steady-state solutions highly depends on parameter 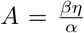, which described the strength of interaction between the morphogen and the curvature, see Figure 3.1. In the weak interplay regime, *A <* 1, we proved the existence of periodic orbits that live completely on the invariant manifold and in the strong interplay regime, *A >* 1, we proved the existence of 2-front periodic orbits; orbits that jump between two invariant manifolds. These findings have biological relevance; it shows that the same system can show different types of patterns depending on the interaction strength between the morphogen and the mechanical response.

The geometric consistency conditions derived in Section 3.1 show that the existence of an orbit in steady-state phase space (3.3) is insufficient for this orbit to be observable as a solution of the PDE. The solution of the ODE system provides the curvature as a function of *s*, which must in addition describe a height function that adheres to the periodic boundary conditions of the curve. A priori, one would expect that these consistency conditions would select a finite number of periodic orbit from a continuous family of steady state periodic solutions. However, by checking one of the conditions, (3.17), we found that a particular orbit is not observable for *ε, δ*_1,2_ too small, see Lemma 3.2/3.5. While *ε* needs to be sufficiently small for the existence proof of periodic solutions (see Theorem 3.1/3.4), the leading order calculation based on the geometric consistency conditions shows that, fixing all parameters, any orbit can be made to violate these conditions when *ε* is sufficiently reduced.

The observability of the periodic solutions was further investigated using numerical simulation; see Section 4. In this section, we vary diffusivity *ε*^2^, curve length *C*, interaction parameter *A* and deviation of curvature-dependent morphogen production from a piecewise continuous function, *δ*_1,2_. The results of the numerical simulations indicate that, while the singular skeleton gives a good approximation of the observed periodic pattern, small deviations from the singular skeleton (either through nonzero *ε* or through nonzero *δ*_1,2_) are necessary to obey the geometric consistency conditions. Furthermore, the simulations in Section 4 show how steady-state solutions change for different parameters. We see that when we increase parameter *A*, the Lagrange multiplier *λ* also increases, as expected from the first order approximation of the integral over the curvature, see Figure 3.3 and 3.10.

The study presented in this paper can provide a framework and inspiration for future research. Many avenues for analytical exploration of model (1.4) are open, which can be used to obtain additional insights into the properties of pattern solutions to (1.4). To bridge the gap between the numerical simulations in Section 4 and the analytical results in Section 3 regarding observability, it would be interesting to include the 𝒪 (*δ*_1,2_, *ε*) correction in the computation of (3.17), as explained in Section 3.5. Moreover, investigating the other geometric consistency conditions from Section might provide extra insights on the observability of the patterns. Furthermore, the stability of these orbits has not yet been analytically tested and could serve as an excellent follow-up research question.

The analysis of periodic patterns in the current paper has produced important follow-up questions by a height function with periodic boundary conditions. Alternatively, one could directly describe a closed curve through its embedding in ℝ ^2^, which could be beneficial for the adherence to the associated geometric observability constraints. One could also extend the current model (1.4), by investigating, for example, the influence of the second term in equation (2.21). Furthermore, the model (1.4) can be altered by taking different functions of *f* (*K*) and the preferred local curvature, *K*_0_(Φ) or making the bending modulus *κ* dependent on the morphogen Φ.

Another important next step would be to apply the results and insights from the analysis presented in this paper to a specific morphogenetic process. The identification of molecules that respond to, or induce, cell curvature [27] can serve as a starting point. Possible applications on tissue scales include Hydra [28] or lung epithelial tissue [17].

## Acknowledgments

The authors would like to thank Alexey Kazarnikov, Moritz Mercker and Anna Marciniak-Czochra for the fruitful discussions and help with the code.

## Notes

https://github.com/DaphnN/Mechanochemicalmodel

### Competing Interest Statement

The authors have declared no competing interest.

